# The Cryo-EM structure of AAV2 Rep68 in complex with ssDNA reveals a malleable AAA^+^ machine that can switch between oligomeric states

**DOI:** 10.1101/2020.06.02.126235

**Authors:** Vishaka Santosh, Faik Musayev, Rahul Jaiswal, Francisco Zárate-Pérez, Bram Vandewinkel, Caroline Dierckx, Molly Endicott, Kamyar Sharifi, Kelly Dryden, Els Henckaerts, Carlos R. Escalante

**Author notes:** To whom correspondence should be addressed: Carlos R. Escalante **Email:**.

## Abstract

The adeno-associated virus (AAV) Rep proteins use a unique AAA^+^ domain to catalyze DNA replication, transcription regulation, and genome packaging. Also, they mediate site-specific integration during a latent phase. To understand the mechanisms underlying AAV Rep function, we investigated the cryo-EM and X-ray structures of Rep68-ssDNA complexes. Surprisingly, Rep68 generates hybrid ring structures where the Origin-Binding-Domain (OBD) forms octameric rings while the helicase domain (HD) forms heptamers. Moreover, the binding to ATPγS promotes a large conformational change in the entire AAA^+^ domain that leads the HD to form both heptamer and hexamers. The HD oligomerization is driven by an interdomain linker region that acts as a latch to ‘catch’ the neighboring HD subunit and is flexible enough to permit the formation of different stoichiometric ring structures. Overall, our studies show the structural basis of AAV Rep’s structural flexibility required to fulfill its multifunctional role during the AAV life cycle.

## Introduction

The adeno-associated virus (AAV) non-structural Rep proteins drive a wide variety of DNA transactions required for the AAV life cycle including, DNA replication, transcription regulation, site-specific integration, and genome packaging (1–7). Remarkably, all these processes are carried out primarily by two functional domains shared by four Rep proteins. The two large Rep proteins (LReps) Rep78 and Rep68, participate in all AAV DNA transactions while the only attributed biological role for the two small Rep proteins (sReps), Rep52 and Rep40, is during genome packaging where they play a central role as the molecular motors (8). The LReps share two domains: an N-terminal origin binding domain (OBD) and an SF3 helicase domain (HD) joined by a linker of ~25 residues (Figure 1A); additionally, Rep78 has a Zn-finger domain at the C-terminus that is not involved in DNA binding (4, 6, 9–11). The sReps consist mainly of the helicase domain, with Rep52 also containing the putative Zn-finger domain (4). As members of the SF3 family of helicases, all AAV Rep proteins have a modified version of the AAA^+^ ATPase domain that lacks the classical C-terminal sensor 2 domain but instead has an N-terminal helical bundle known as oligomerization domain (OD) (12, 13). X-ray and biochemical studies have shown that the OBD has three non-overlapping DNA binding motifs that carry out distinct functions: A HUH-fold catalyzes the cleavage of ssDNA in the terminal resolution reaction during DNA replication (14–16); a dsDNA binding pocket that recognizes Rep binding sites (RBS) found in the origin of replication, p5 promoter site and AAVS1 integration site (11, 15, 17); and a third motif that binds ssDNA hairpins (14, 15, 18). The helicase domain has an additional DNA binding motif known as the pre-sensor 1 β-hairpin (PS1βH), common in all SF3 helicases and involved in melting and unwinding (12, 19). Taken together, the LReps have multiple DNA binding elements that can potentially alternate or cooperate to catalyze all AAV DNA transactions.

**Figure 1.**
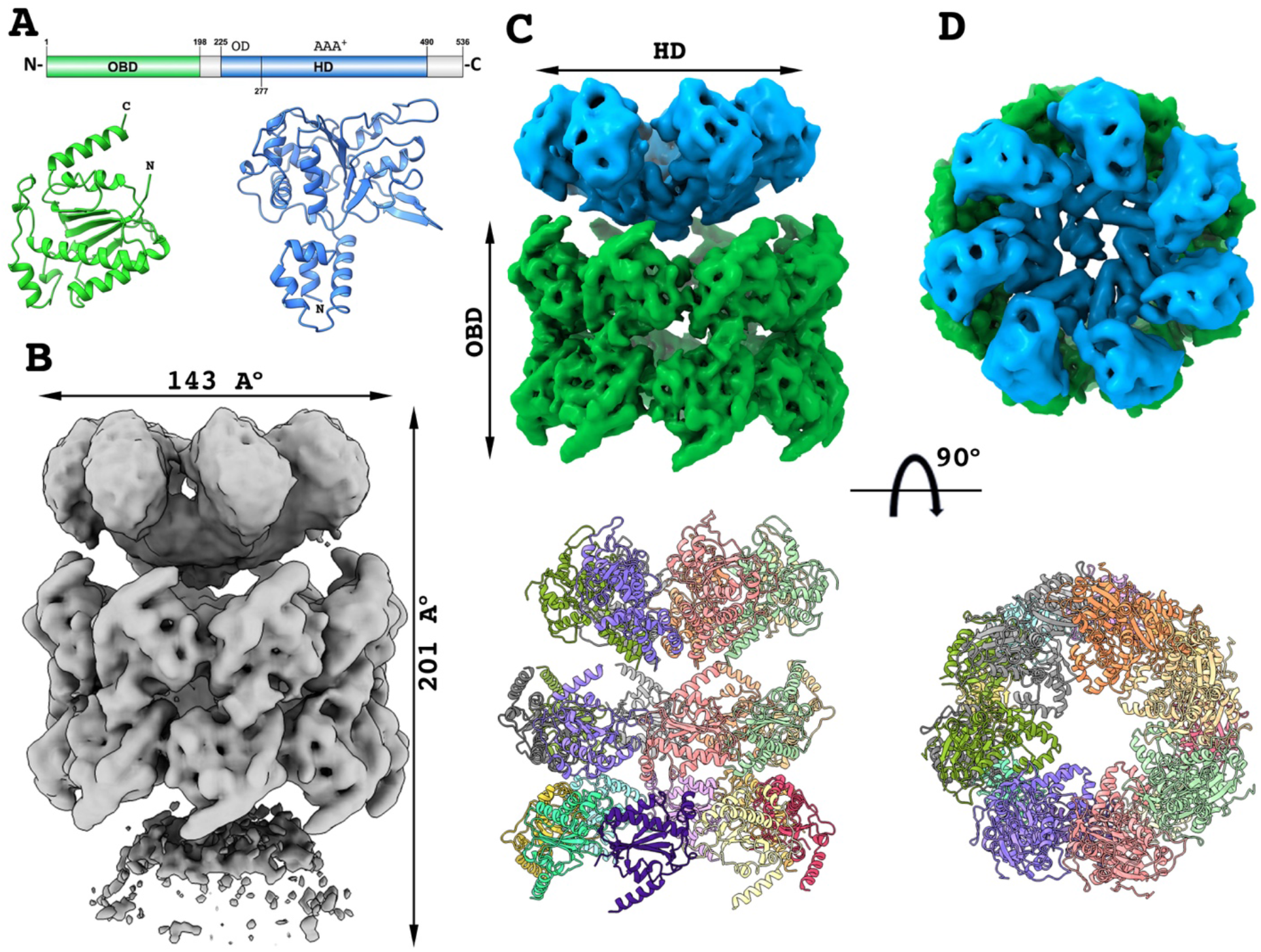
Overview pf the Rep68-ssDNA DOC. (A) Primary and X-ray structures of Rep68 functional domains. Color scheme is maintained throughout all figures. OBD: origin binding domain; HD: helicase domain; OD: oligomerization domain. (B) 3D Cryo-EM map of the Rep68-ssDNA DOC and overall dimensions. (C-D) Top panel: Composite local cryo-EM maps of HD1(blue) and OBD DOC (green), Bottom panel: Ribbon representation of the corresponding atomic models. X-ray structures of AAV-2 OBD (PDB:5BYG) and AAV-2 Rep40 (PDB: 1S9H) were used to generate all atomic models.

Although the overall domain structure of the AAV Rep proteins is similar to those of other viral SF3 helicases, they could be considered a subclass of this helicase family because of their distinctive oligomerization properties. Whereas common viral SF3 helicases and most AAA^+^ proteins form only hexameric rings, Rep40 -containing only the helicase domain-is a monomer in solution and only forms transient dimers in the presence of ATP (16, 20, 21). In contrast, the LReps display a dynamic association process forming a spectrum of oligomeric species, from dimers to octameric rings (20). Furthermore, studies show that the interdomain linker plays an active role in the oligomerization of the LReps as its substitution by a non-related linker, completely abolishes the ability of Rep68 to oligomerize (22). Therefore, the minimum AAV Rep helicase domain -as represented by Rep40-is deficient in its ability to oligomerize while oligomerization in the LReps requires the cooperative interaction of multiple domains producing complexes that are intrinsically dynamic. More importantly, this conformational plasticity allows LReps to acquire different quaternary structures modulated by the nature of the DNA site. For instance, binding to the integration site AAVS1 produces a heptameric complex, whereas binding to poly-dT ssDNA results in the formation of a double-octamer (23). To understand the structural basis of these processes, we determined cryo-EM structures of Rep68 in complex with ssDNA. The structures reveal that each functional domain can form oligomers with different stoichiometries. Thus, while the OBD favors the formation of octamers, the HDs forms heptamers. Upon binding

ATPγS, the HDs transition into hexameric rings. Moreover, the structures show that the key structural motif inducing HD oligomerization is a linker region of 11 residues that acts as a flexible latch docking into neighboring OD domains and stabilizing both heptamers and hexamers. Our results provide essential information that explains the remarkable conformational plasticity of AAV Rep proteins that makes them a unique subclass of SF3 helicases, illustrate the versatility of the AAA^+^ domain, and offer insights into their DNA remodeling functions.

## Results

### Overall architecture of the Rep68-ssDNA complex

To gain insights into the molecular mechanisms underlying the multifunctionality of AAV Rep proteins and their unique oligomeric properties, we carried out a combination of cryo-EM and X-ray studies of several Rep68-ssDNA complexes. First, we focused on our previously characterized Rep68-ssDNA double-octameric complex, from here on referred to as DOC (23). Using single-particle cryo-electron microscopy, we were able to obtain a cryo-EM map at an average resolution of 4.6 Å (Figure 1B, Supplementary Figure S1, Table S1)). The structure of the complex resembles a double funnel with overall dimensions of 201 Å by 133 Å by 143 Å (H × W × D). The map can be divided into three ring-like sections: a well-defined central ring composed of OBDs (RING_OBD_) flanked by two outer helicase rings (HD1, HD2). The density of HD2 is too weak and fragmented to identify individual subunits (Figure 1B). Likewise, the densities of the linkers connecting the OBDs to the HDs are not visible, indicating that they are highly dynamic. A close examination of the RING_OBD_ map shows that it consists of two octameric OBD rings arranged in a head-to-head orientation and related by pseudo-D8 symmetry (Figure 1C). Surprisingly, there is a symmetry mismatch between the OBD and HD rings, with HD1 showing seven well-defined domains while the HD2 density is too diffuse to assign a stoichiometry (Figure 1C-D). We could detect a “satellite” density on the periphery of the HD1 ring, which could represent the missing eighth domain, but it could not be resolved well enough to confirm this observation. Altogether, the results show that the DOC has a central stable core of two octameric OBD rings and two highly dynamic HDs rings.

### The double-octamer is stabilized by ssDNA

The OBD double ring is 94 Å wide and with an outer diameter of 143 Å generating an internal channel with a diameter of ~ 67 Å (Figure 2A). The cryo-EM map of the RING_OBD_ is of good enough resolution to fit the complete main-chain backbone from the OBD X-ray structure and some of the bulkier side chains (residues 1-198) (Figure 2A). Sixteen individual OBDs can be easily docked to generate a double-octameric OBD ring model. The only region from the X-ray structure that does not fit entirely into the cryo-EM density is the C-terminal α-helix F that is slightly tilted by 2° with respect to the cryo-EM density but was fitted into the density using rigid body refinement. The OBD molecules are oriented such that the HUH catalytic residues reside in the inside of the ring. Surprisingly, the interface involved in generating the octameric ring utilizes the same motif that recognizes the dsDNA RBS GCTC repeats, namely, helices D and the loop L_DB_ (Figure 2B)(17). This region interacts with α-helices B, C, and the loop connecting strands β2 and β3 in the neighboring OBD molecule (Figure 2B). In all, this interface potentially buries 1180.5 Å^2^ of solvent accessible area for a total of ~4720 Å^2^/octamer (17).

**Figure 2.**
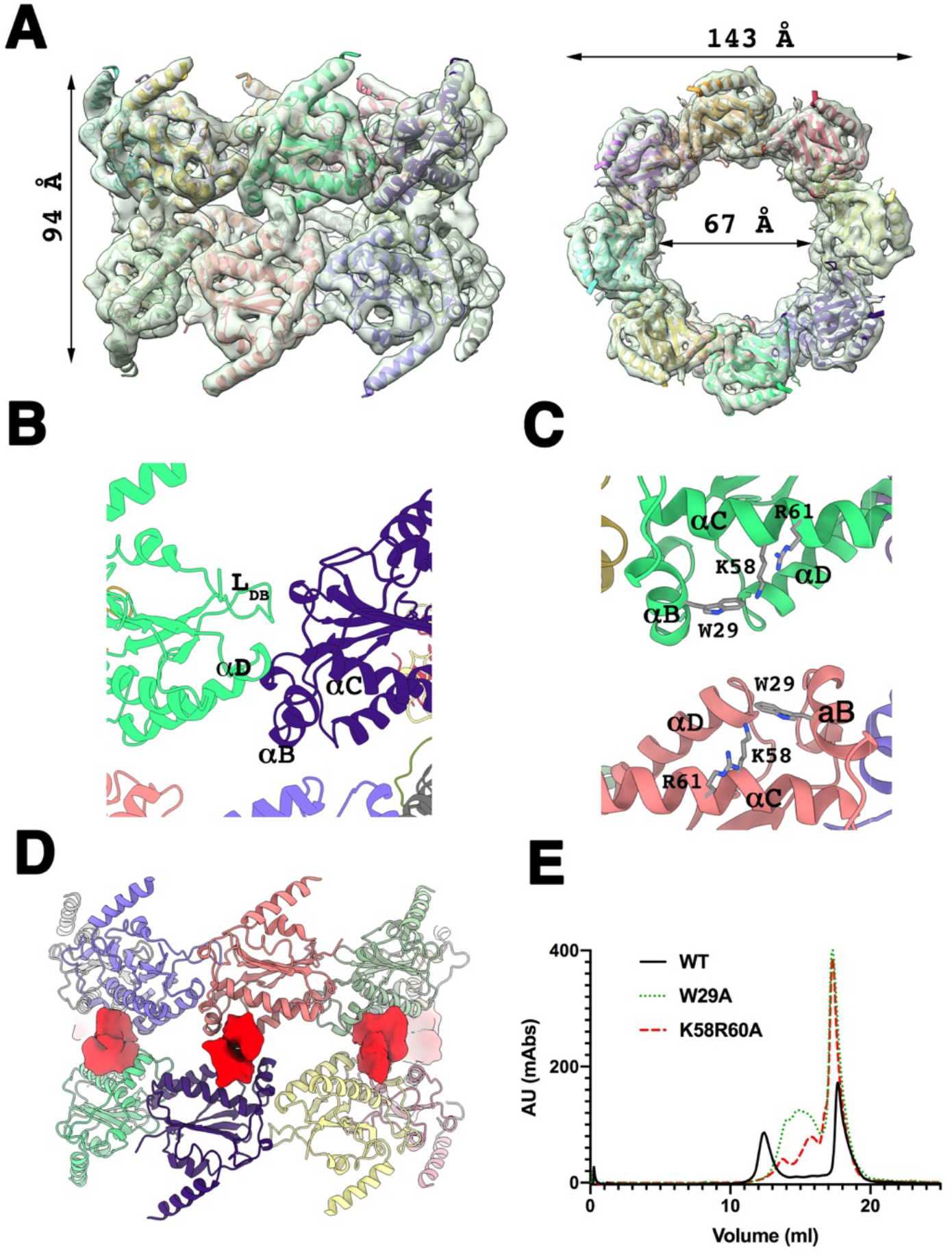
Structure of OBD Double-octamer. (A) Two orthogonal views of the local Cryo-EM maps and 16 refined OBDs. (B) Ribbon representation of the OBD-OBD interface in the octamer. The L_DB_ (DNA binding loop) and αD interact with αB and αC residues in the neighboring subunit. Both motifs also recognize the GCTC repeats in RBS DNA sites. (C) Ribbon representation of the interface generating the DOC. Each OBD is related by D8 symmetry. Three side chains W29, K58 and R61 have been identified as critical for ssDNA binding. (D) Unaccounted density (red) located between OBD molecules in the crevice found between αB-C and αD. (E) Size exclusion chromatography profiles of W29A and K58R61A mutants and their ability to form the DOC. Samples were run on a Superose 6 column, with Rep68 WT-ssDNA DOC eluting at ~12 ml (black line), W29 mutant (green dotted) elutes at 15 ml and K58R61A at 16. ssDNA elutes at 18 ml.

The interactions mediating the formation of the double-octameric ring comprise a pocket generated by α-helices B, C, and D in each of the sixteen OBD molecules (Figure 2C). Interestingly, this region contains three residues (W29, K58, R61) that were previously identified as critical to binding ssDNA hairpin RBE’ found in the AAV5 ITR (15). A closer inspection of the Cryo-EM density around this pocket, shows an unaccounted density that spans neighboring OBDs across individual octameric rings (Figure 2D). We hypothesize that the unaccounted density corresponds to ssDNA molecules bridging OBD rings. To test this hypothesis the predicted interacting residues (W29, K58, and R61) were mutated to alanine and were tested for their ability to form the complex. Figure 2E shows that Rep68 containing any of the mutations fail to form the complex. To understand the DNA structural requirements that lead to the formation of the Rep68 DOC, we carried out size-exclusion chromatography experiments with different substrates. First, we aimed to identify the minimal length of ssDNA required to form the complex using ssDNA molecules of ten, fifteen, seventeen, and twenty-five thymines. The elution profiles of each sample show that ssDNA smaller than 17 nucleotides failed to form the complex (Supplementary Figure S2A). Next, we explore the effect of the composition of the ssDNA sequence using poly-guanine, poly-cytosine, and a region from the integration site AAVS1. Elution profiles show that the complex forms more readily using ssDNA molecules that are pyrimidine-rich, with preference dT>dC>dG (Supplementary Fig. S2B).

### The X-ray structure of OBD with ssDNA shows formation of double-octameric rings

In previous work, we showed that Rep68 forms octameric rings, which represent one among an assortment of oligomers present in solution (20). Based on this data, we hypothesized that the OBD by itself may assemble as an octameric ring under the right conditions, mainly at high enough concentrations. To test this hypothesis, we performed sedimentation velocity experiments at different concentrations and compared the experimental sedimentation coefficient to the sedimentation coefficient calculated for different OBD atomic models, including an OBD octameric ring using the program SOMO (24, 25). Results show that at concentrations below 124 μM, the OBD forms two different species, one sedimenting at ~ 2.2 S and a faster species at ~5 S (Supplementary Figure S3A). Higher concentrations result in additional species with a sedimentation coefficient of 7.2 S, which is similar to that predicted for the octameric ring (7.3 S) (Supplementary Figure S3A). In addition, we also performed glutaraldehyde crosslinking of OBD, showing the formation of octamers on an SDS-PAGE (Supplementary Figure S3B). These experiments demonstrate that the formation of octameric rings is an intrinsic property of the OBD.

Furthermore, we determined the crystal structure of OBD in complex with a ssDNA molecule containing an RBS sequence from the AAV2 ITR. The complex crystallizes in space group I222 with four OBD molecules in the asymmetric unit sharing two molecules of ssDNA (Table S2). The arrangement of the four OBD molecules resembles that of the cryo-EM Rep68-DOC using the same OBD-OBD interface motifs with ssDNA bridging two molecules (Figure 3A). Moreover, analysis of the arrangement of the molecules in the crystal lattice shows that the OBD assembles as double octameric rings (Figure 3B). The dimensions of this ring are identical to the cryo-EM structure such that a model of the octameric ring from the crystal structure can be manually docked into the cryo-EM density producing an almost perfect fit. We can safely predict that the interface residues we observe in the crystal structure are the same as what we observe in the cryo-EM structure.

**Figure 3.**
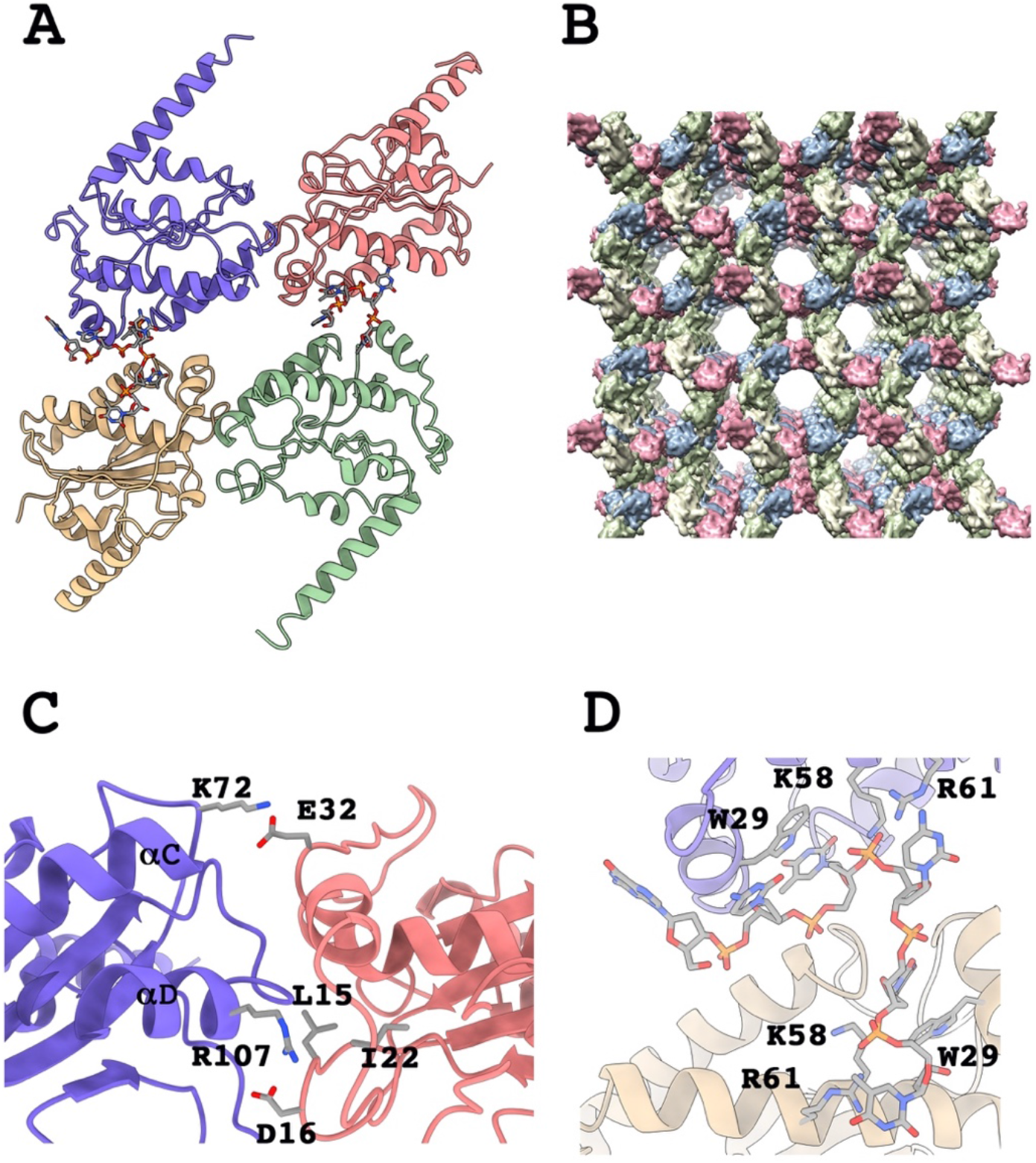
X-ray Structure of OBD-ssDNA complex. (A) Overall view of the asymmetric unit with four OBD molecules bridged by two ssDNAs. (B) A view of the crystal lattice showing the double octameric assembly. (C) Ribbon representation of the OBD-OBD octameric interface and some of the most representative residues involved in protein-protein interactions. (D) Structure of the bridging ssDNA showing stacking interactions of W29 with thymines and K58 and R61 interacting with the phosphate backbones.

Examining the protein contacts, we were able to identify the specific OBD-OBD interface residues forming the octameric ring. An example of such interactions occurs between α-helix C residue K72, forming a salt bridge with E32 on helix B of the neighboring OBD molecule (Figure 3C). Of particular interest is residue R107 in helix D that forms multiple interactions with L15, D16, I22, and S23 of the neighboring OBD molecule (Figure 3C). Not surprisingly, an R107A mutation is defective in Rep68*-ssDNA complex formation, as we had previously shown in the initial Rep68-DOC characterization study (23).

The crystal structure also shows how the DOC assembles by the ssDNA bridging of two octameric rings. The DNA used in the crystallization was a 14-mer with sequence 5’-ATATGCTCG<u>CTCTT</u>-3’ that adopts a stem-loop conformation. A pair of OBD molecules bind to the 3’-half of the ssDNA molecule interacting with the last five nucleotides (CTCTT). The sequence TCT interacts across two OBD molecules with each thymine making stacking interactions with residue W29 and forming a cation-π interaction with R122 in each OBD (Figure 3D). Other key residues from this region are K58 and R61 that make phosphate backbone interactions with the middle cytosine of this tri-nucleotide region. This interface is equivalent to the one found with AAV5 OBD in complex with the RBS’ stem-loop and suggests that the OBD ssDNA binding motif has a strong preference for pyrimidine rich DNA (15). Moreover, it explains, as previously noted, how mutations of residues W29, R58, and K61 abolish the formation of the Rep68-DOC.

### Heptamerization of the helicase domain is directed by the OD

The Rep68 HD1 region shows seven well-defined densities forming a two-layered ring of different diameters. The smaller tier ring is made up of the N-terminal oligomerization domains (ODs), forming a continuous ring with an external diameter of 75 Å (Figure 4A). The larger tier with a diameter of 134 Å is discontinuous, with well-separated densities consisting of 7 AAA^+^ domains (Figure 4A). The resolution of the ODs is similar to that of the OBDs and we can trace the complete main chain (residues 224-277). In contrast, the resolution of the AAA^+^ domain is significantly lower, due to its inherent dynamics. 3D variability analysis shows that the HD1 ring is very dynamic, moving as a rigid body with respect to the core OBD octamer, and also each individual helicase domain moving independently, particularly the AAA^+^ subdomain (Supplementary Figure S4). The overall HD1 ring has a funnel-like inner channel with a decreasing radius from 76 Å at the top of the AAA^+^ ring to 25 Å at the lower end of the OD ring (Figure 4B). Surprisingly, the HD1 model shows that heptamerization is only mediated by interactions between the ODs as no interaction between the AAA^+^ domains are detected (Figure 4A). The X-ray Rep40 structure fits without any major changes in its overall conformation. The cryo-EM density also indicates that the N-terminal α-helix 1 in the OD can be extended with 4-6 residues from the linker region (Figure 4C). The OD-OD interface buries 536 Å^2^ of surface area and is made up of helix 1 and 4 in one subunit and helices 2 and 3 in the neighboring OD as previously predicted (26). Although the cryo-EM map is not of sufficient resolution to reveal side chain interactions, the fitted X-ray atomic model reveals multiple potential interface interactions between OD subunits, particularly N254 and S261, that are in close proximity to Y230. To test this potential interface, we mutated both residues to alanine and measured the effect on viral replication in an infectious particle assay. Figure 4D shows that each mutation causes a dramatic decrease in the production of infectious AAV particles implying the predicted interface residues are functionally significant.

**Figure 4.**
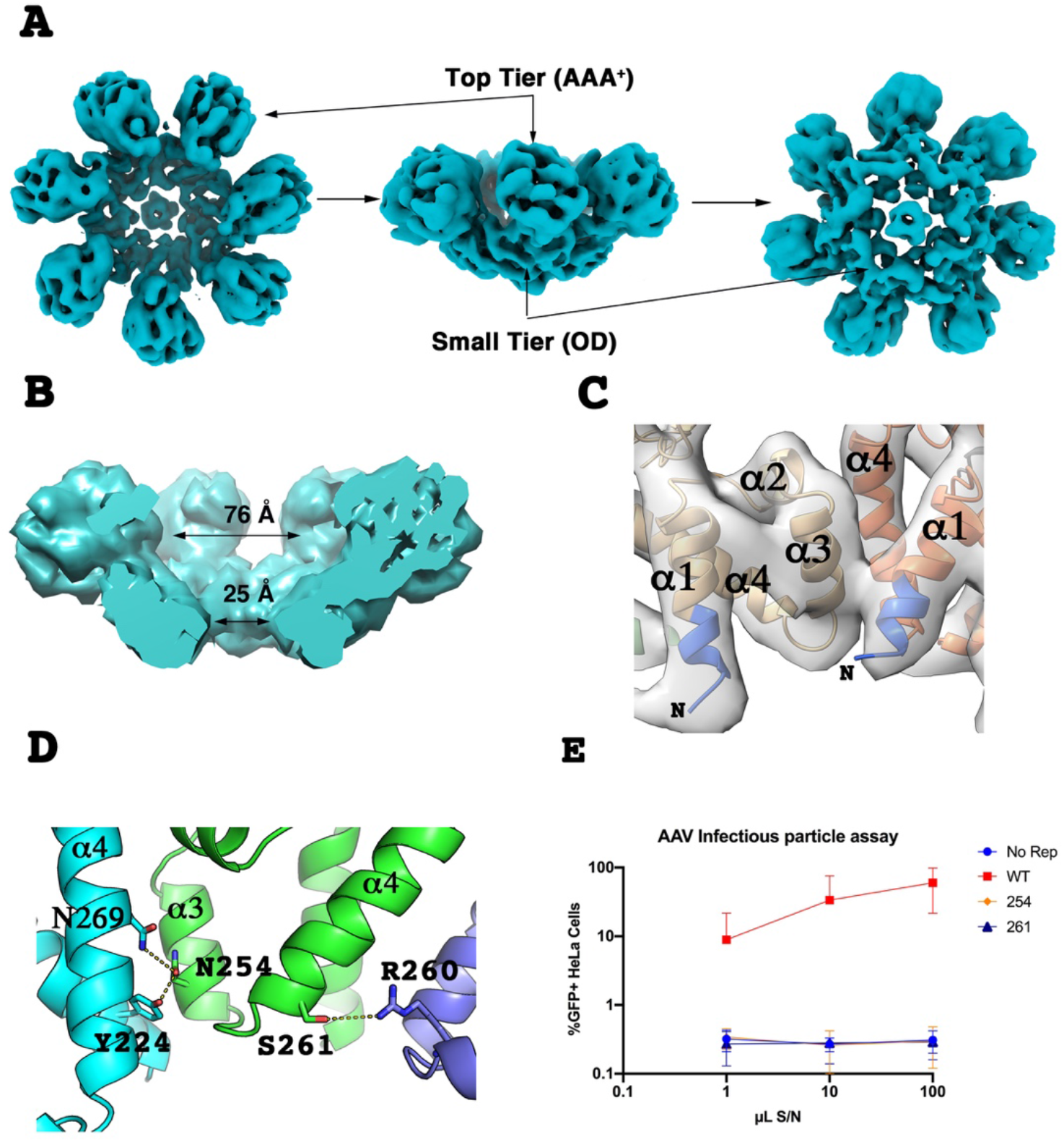
Structure of APO Heptameric Helicase domain. (A) Local Cryo-EM density of top (left), side (middle) and bottom (right) views. (B) Cutoff of the side view of the HD showing the different diameters along the central channel. (C) Cryo-EM map and fitted Rep40 structure around the OD showing additional density that extends the N-terminal helix (in blue). (D) Ribbon representation of some of the interactions involved in the OD-OD interface. The view is from the center of the channel. (E) Infectious particle assay using WT Rep and the N254A and S261A Rep mutants (n=4).

### The OD contains a second DNA interacting motif

There is an additional density at the center of the HD1 heptameric ring structure that likely represents part of a ssDNA molecule (Figure 1A,4A). The size and shape of the density fit a ssDNA model of ~four nucleotides (Figure 5A). Surprisingly, the DNA density is not located around the DNA translocation pre-sensor 1 β-hairpin (PS1βH) but resides instead at the N-terminal end of the OD ring, where several basic residues such as R260 and K264 generate an electropositive surface area (Figure 5B-C). The presence of ssDNA at this location is a reflection of the larger electropositive charge in this region as compared to the PS1βH ring (Figure 5C). To test the role of these residues, we mutated R260 and K264 to alanine and performed an infectious particle assay (26). Figure 5D shows that both mutations completely abolish the formation of infectious AAV particles indicating they play a critical functional role during the AAV life cycle. However, helicase assays show that the mutants are still active, although to a lesser degree than the WT protein, implying they play a role other than supporting helicase function during the life cycle of AAV (Figure 5E). Lastly, because the HD2 does not appear to have an equivalent DNA density in the channel, it may explain its high dynamic character, as the ssDNA provides stability for the association of the OD rings.

**Figure 5.**
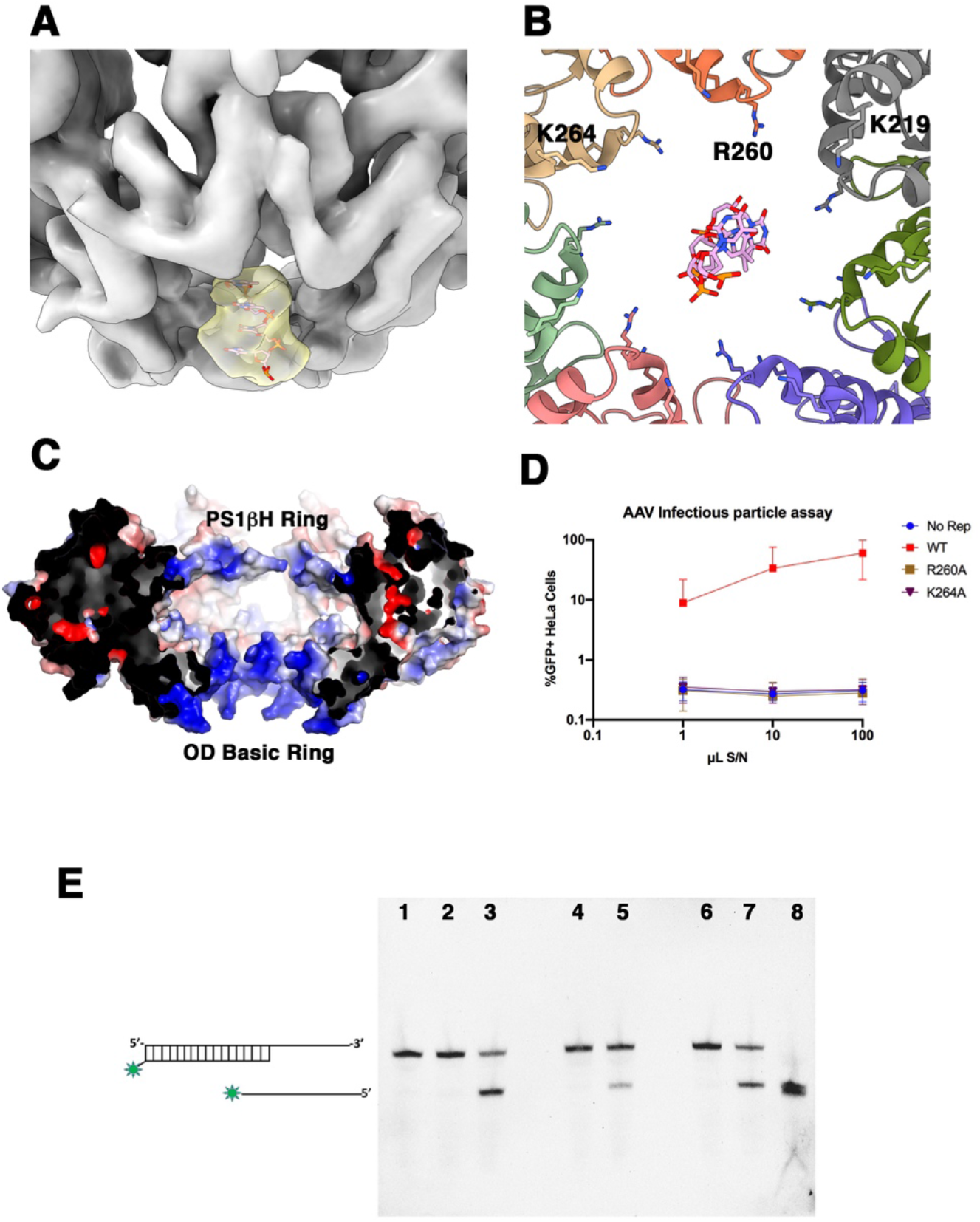
ssDNA molecule in the center of the channel. (A) Side-bottom view of the HD Cryo-EM map displaying DNA density (yellow) and a docked poly-thymine (dT4) model. (B) Bottom view of the lower tier OD ring showing residues R260 and K264. (C) Cutoff showing surface representation of the electrostatics inside the central channel. Two electropositive rings (blue) are located in the AAA^+^ domain at the PS1βH and at the OD. (D) Infectious particle assay of R260A and K264 mutants showing a drastic effect in the production of infectious particles compared to WT (n=4). (E) Helicase assay. Lane 1: fluorescein-dsDNA substrate; lane 2: WT no ATP; lane 3: WT+ATP; lane 4: R260A no ATP; lane 5: R260A + ATP; lane 6: K264A no ATP; lane 7: K264A + ATP; lane 8: fluorescein-ssDNA.

### In the presence of ATPγS, the Rep68 helicase heptamer is in a dynamic transition between open and close rings

The Cryo-EM structure of the ATPγS-bound Rep68-ssDNA DOC complex was solved to an overall resolution of 5.2 Å (Supplementary Figure S5). Interestingly, during 2D classification, hexameric ring particles were also identified. Overall, there are no significant changes in the DOC architecture except in the helicase domain. Although the map density of the helicase domain is less defined than the apo structure, we can still observe that HD1 still forms a heptameric complex. After local refinement of the HD, we fitted the OD and AAA^+^ domains as different rigid bodies as the density showed significant differences when compared to the apo structure (Figure 4A,6A). The diameter at the top of the inner channel constricts to 46 Å generated by an ~44° rigid body rotation of the AAA^+^ domains toward the center of the ring (Figure 6B). There is a density that spans the length of the central channel and protrudes at the lower OD-tier corresponding to ssDNA (Figure 6C). Several subunits are clearly contacting DNA through residues of the OD ring and the AAA^+^ domain, although the resolution is not sufficient to see specific side chains; however, it is clear that residues from the PS1βH are involved (Figure 6D). A combination of factors may account for the lack of definition of the HD1 cryo-EM density, including the presence of multiple conformations and multiple HD stoichiometries. Because the DOC complex may not be the most representative of the functional complexes that LReps form during the viral life cycle, we decided to generate single-ring Rep68 complexes using more physiologically relevant ssDNA sites to study the changes induced by nucleotide binding to the helicase domain. This was achieved by purifying a Rep68-ssDNA-ATPγS complex containing a fifteen-nucleotide sequence from the AAVS1 integration site. 2D class averages show a mixture of HD heptameric and hexameric rings as in the DOC-ATPγS complex (Figure 7A). It is clear that the formation of hexameric rings is nucleotide-dependent because they are only present in samples that contain ATPγS. The 2D classes show no density for the OBDs, suggesting they are highly mobile once they are not engaged in forming octameric rings. The new HD heptameric complex was solved to 4.6 Å, a similar resolution as the apo complex (Supplementary Figure S6). The cryo-EM map shows density for ssDNA spanning most of the inner channel from PS1βH to the OD domain (Figure 7B). More notably, the ssDNA density shows a clear curvature suggesting DNA deformation occurs. The seven individual AAA^+^ subunits are distinguishable, but the OD domains are less defined, forming a continuous density that is incomplete in some regions, suggesting a very dynamic complex (Figure 7B). We undertook a *3D* variability analysis using cryosparc v2.15 to determine the conformational heterogeneity of the heptameric complex (27, 28). Results show that the heptamer exists in at least three distinct conformations with maps showing open and closed rings and also rings divided into halves of four and three subunits (Figure 7C). Taken together, the ATPγS-bound structures show that Rep68 undergoes a series of conformational changes that induce assembly-disassembly of the heptamer and potential DNA deformation.

**Figure 6.**
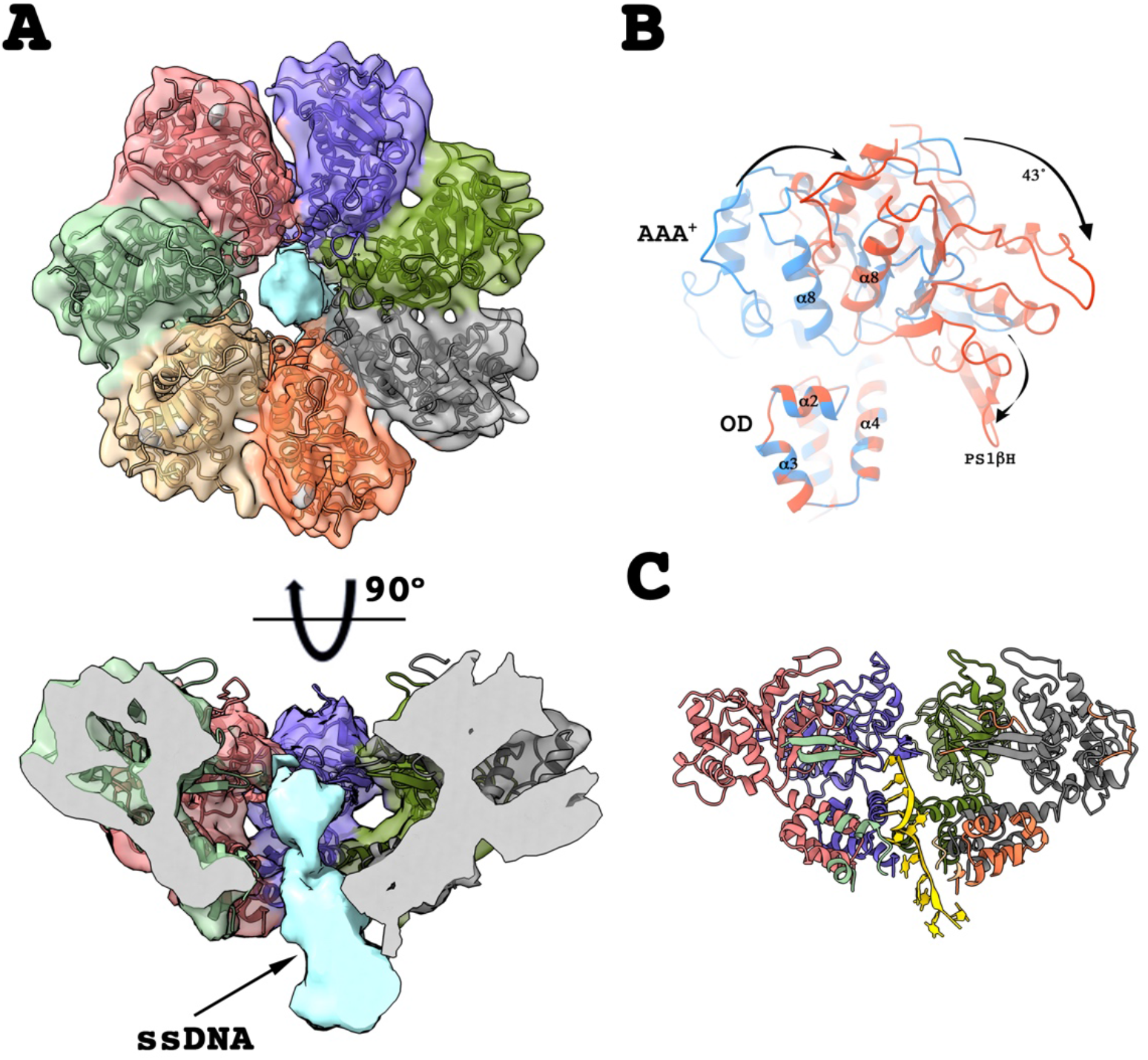
Effect of ATPγS on DOC complex. (A) Top view of HD density and fitted heptameric model, blue density in the central channel is ssDNA. (B) Superposition of APO (blue) and ATPγS (red) helicase domains showing the large conformational change of the AAA^+^ domain. The HDs were superposed using only the ODs. (C) Cutoff of a side view of the heptameric complex showing the ssDNA spanning most of the channel (left) and a ribbon representation of the model (right).

**Figure 7.**
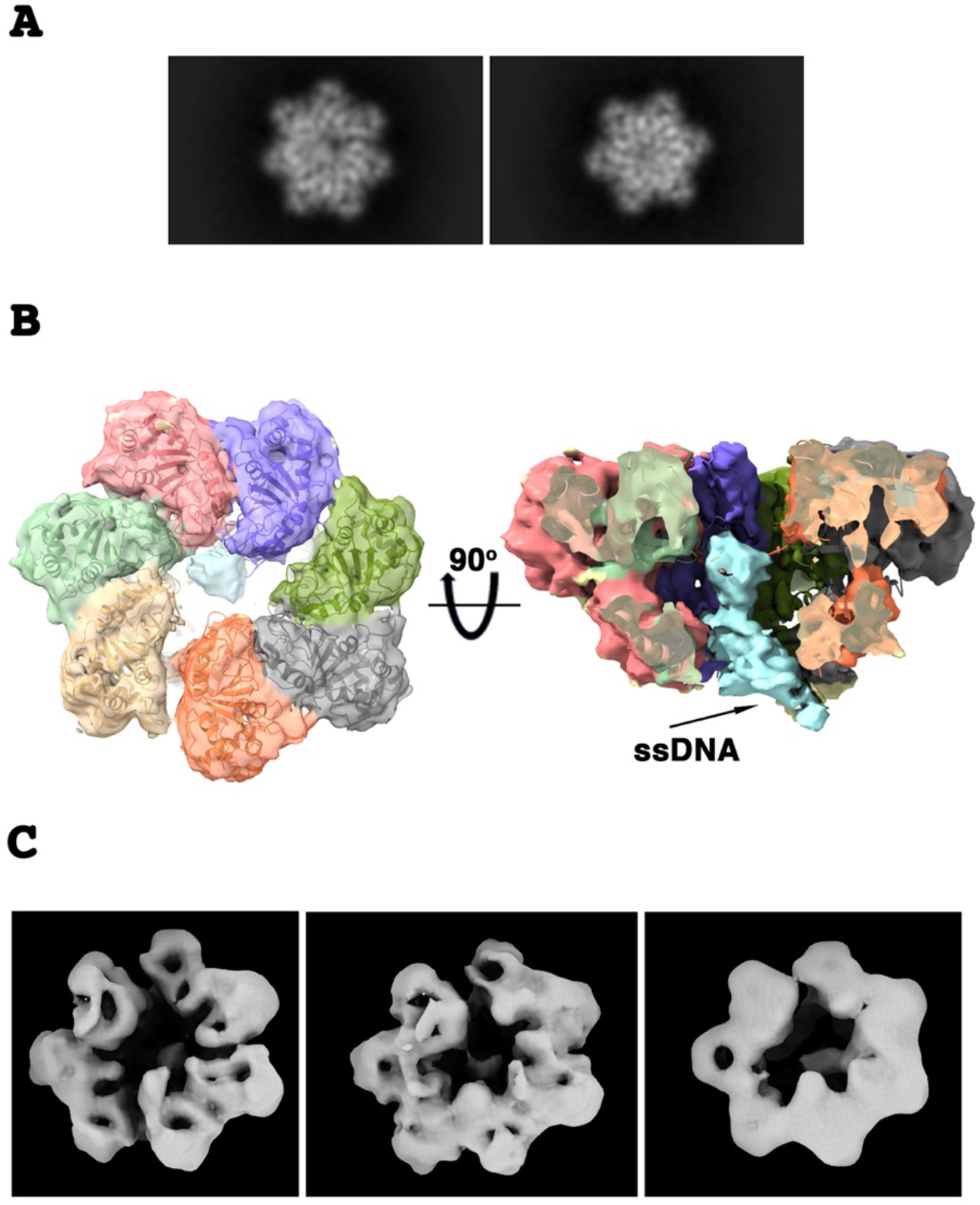
Cryo-EM structure of Rep68-ssAAVS1-ATPγS heptameric complex. (A) Representative 2D classes showing heptameric and hexameric ring particles. (B) Top (left) and side (right) views of the Cryo-EM density with fitted helicase model. Blue density in the central channel represents ssDNA. There is a clear induced curvature of the DNA molecule. (C) Representative Cryo-EM maps of three conformations of the Rep68-ssAAVS1 complex found through 3D variability analysis.

### The hexameric complex shows the linker forms a flexible latch that stabilizes the oligomeric rings

As previously mentioned, 2D classification of ATPγS complexes detected hexameric ring particles with both poly-dT and AAVS1 ssDNA. We determined the cryo-EM structure of the Rep68-dT25 hexameric ring complex as the ssAAVS1 complex did not have enough particle orientations to generate a high-resolution model. The complex was solved to an overall resolution of 5.0 Å after local refinement (Figure 8A, Supplementary Figure 7). The overall arrangement of the hexamer resembles that of other SF3 family members, but with significant differences. Particularly, the arrangement of the subunits does not follow a proper 6-fold rotational symmetry, displaying differences in how each subunit docks into the ring. The diameter of the ring is 116 Å in width and 69 Å in height with an inner tunnel that has narrowed to ~ 13 Å. The rigid body rotation of the AAA^+^ domain with respect to the apo structure is ~40°, similar to the heptameric ATPγS complex. Most of the density for ssDNA resides along the region of the PS1βH region with some minor density around the entrance of the OD. The Cryo-EM map shows additional unaccounted density that extends the N-terminal end of the OD to include five additional residues from the linker region. Part of this region docks into a crevice formed between OD helices 1 and 3 of the neighboring molecule in a clockwise direction (Figure 8B-C). Residues V215 and I216 in the linker most likely form part of a hydrophobic pocket that includes W230 from helix 1 (Figure 8B). Looking back at the heptameric apo HD1 structure, we can also now distinguish density for the latch region that wasn’t accounted for before due to the lower resolution of the map Figure 8C). Consequently, the latch motif is flexible enough to stabilize ring structures with different stoichiometries producing rings that have a dynamic assembly/disassembly process. This property is illustrated by the variety of conformations that we see in the heptameric ring (Figure 7) and the 3D variability analysis of the hexameric ring. Results from this analysis show that in one of the normal modes, a subunit coming in/out of the ring (Figure 8D, SM1). Moreover, the 2D classes show a minor population of particles with only five subunits (Figure 8E). Taken together, our results imply that upon ATP binding and/or hydrolysis, there is formation of hexameric rings, possibly by disassembly of heptamers.

**Figure 8.**
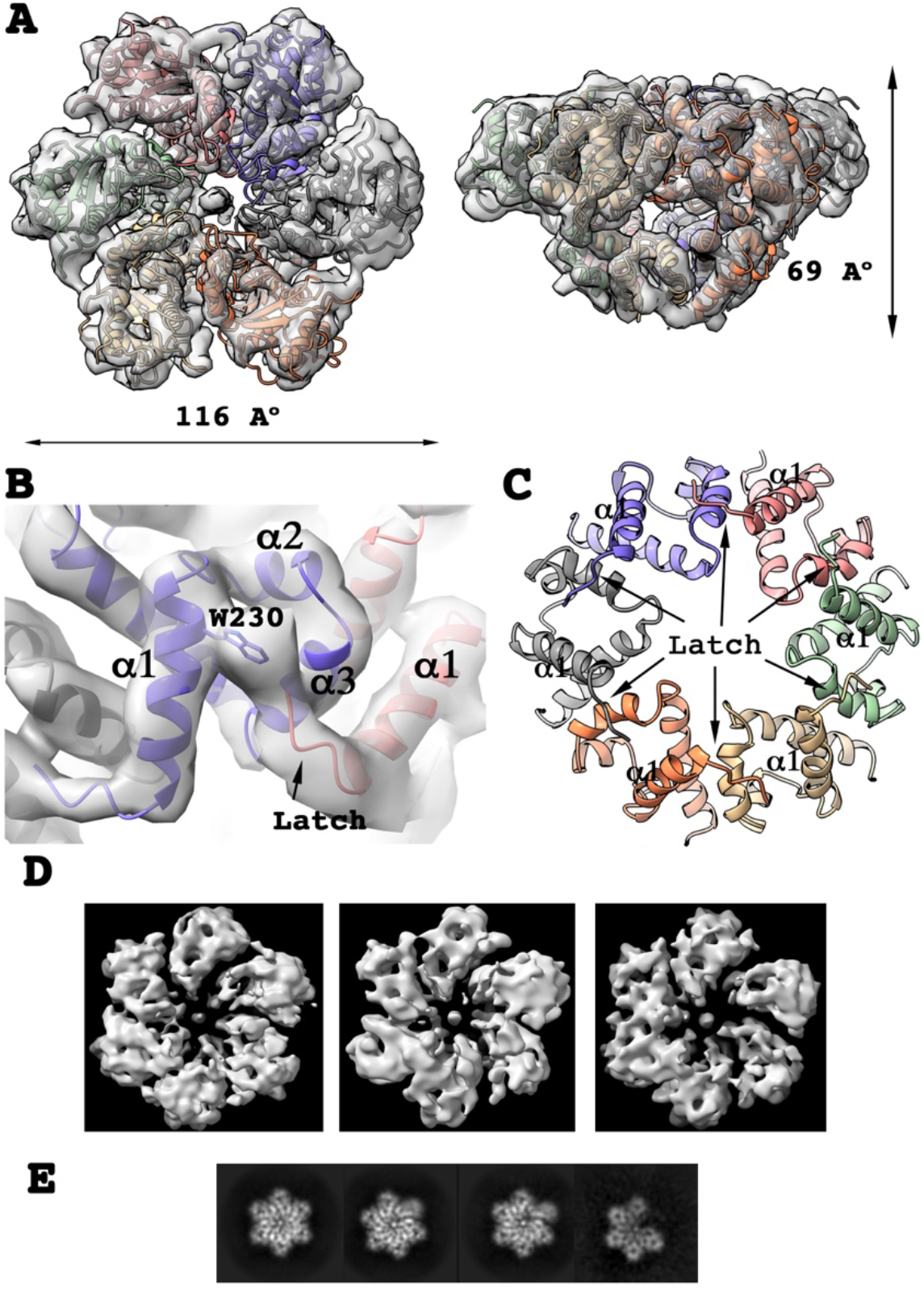
Cryo-EM structure of Rep68-ssAAVS1-ATPγS hexameric complex. (A) Top and side views of the cryo-EM hexameric complex with fitted helicase model. (B) Details of the flexible latch (pink) docking into a pocket of the neighboring subunit. (C) Ribbon representation of the ODs with and the latch of each subunit in both heptameric and hexameric rings. (D) Cryo-EM maps of three normal models obtained through 3D variability analysis. There is an in/out movement of one of the subunits. (E) Representative 2D classes of the hexamer and its dynamic behavior. The last class shows a five membered open ring.

## Discussion

The cryo-EM and X-ray structures of AAV Rep68 in complex with ssDNA illustrate the structural basis that confers AAV Rep proteins the conformational plasticity necessary to acquire multiple oligomeric states upon binding to different types of DNA substrates. This oligomeric malleability originates from three key features: the presence of multiple oligomerization interfaces located in three different domains, the existence of several protein-DNA interaction motifs and the fact that each domain can form different stoichiometric oligomers independent of each other, resulting in the OBDs associating as octamers and the HDs forming heptamers and hexamers. The active participation of the OBD in oligomerization was unexpected, as no reports from other members of the SF3 and HUH families having this property have been described previously. As shown by our AUC and X-ray studies, the propensity to form octameric rings is inherent to the OBD, as it can form octamers at high concentrations (> 200 μM), illustrating that this oligomerization interface is not particularly robust. However, in the context of the LReps and under the right conditions, the formation of octamers occurs naturally (20). It is clear from the cryo-EM and X-ray structures that the formation of the double-octameric complex occurs through ssDNA bridging of two octameric complexes requiring a ssDNA molecule with a minimum size of 17 nucleotides in length and containing a stretch of at least three pyrimidines, preferably thymines. At the present time, we don’t know if the DOC participates in any of the multiple processes during the life cycle of the AAV. However, a simple inspection of the AAV genome sequence shows that sections with three and four consecutive thymines occur at least 50 and 12 times, respectively, and segments with successive thymines and cytosines at higher frequencies. This suggests that, in principle, there are enough sites in the AAV genome for the formation of the double-octameric complex. Still, it is also possible that the DOC-structure only forms in vitro as a result of the conditions used in our studies, particularly the use of a dT_25_ ssDNA. Nevertheless, the use of poly-dT has permitted us to trap the Rep68 octameric complex in a conformation that may be difficult to capture due to the inherently dynamic nature of the system. This dynamic character can be seen in the cryo-EM structures of the heptameric Rep68 with ssAAVS1 (Figure 7) and Rep68-dT_25_ hexamer (Figure 8), where the OBD domains by being not engaged in octameric interface formation, are not resolvable due to their high mobility.

The question remains as to what the functional significance of the OBD octameric structure is. One of the most intriguing features of the DOC is that the interface utilizes the same motifs -and some of the same residues-as those that participate in the specific recognition of RBS dsDNA sequences found at the AAV origin of replication, p5 promoter, and AAVS1 integration site. Therefore, it is tempting to postulate that after the initial binding of Rep to RBS sites, the interface switches from a DNA recognition mode into an oligomerization mode. This arrangement puts the OBD ring in a conformation such that the HUH nuclease catalytic core is located in the exterior face of the ring, thus preventing any unintended hydrolysis of the DNA backbone. Further studies will be needed to explore this hypothesis.

We had previously shown that the interdomain linker plays an active part in Rep68 oligomerization; the work presented here provides the structural determinants of such function (22). The cryo-EM structures illustrate that as predicted, the linker is partly folded and extends the N-terminal helix of the HD by ~ 6 residues (18, 22). More importantly, five additional linker residues preceding this N-terminal helix form a flexible latch that docks into the OD of a neighboring subunit (Figure 8C). Remarkably, this same interaction can be observed in both the heptameric and the hexameric complexes. We propose that this region (aa 215-220) acts as a flexible latch to induce oligomerization and stabilize the ring structures. Is this motif unique to Rep proteins? Analysis of the X-ray structures and biochemical studies of SV40-Tag show that it assembles as hexamers without the need of a linker latch region (29, 30). Similarly, the first E1 helicase X-ray structure lacked the equivalent linker residues but still was able to form hexameric rings (31). However, subsequent structures with E1 constructs that included the linker region show a similar flexible latch that interacts with the neighboring subunit, although in opposite direction (32). Thus, although the latch motif may not promote oligomerization in other SF3 helicases, it is absolutely necessary in AAV Rep proteins and can be considered an integral part of the helicase domain.

Another unexpected feature of the apo Rep68 structure, when compared to other SF3 family members, is that the AAA^+^ domain does not participate in the oligomerization process (Supplementary Figure S8). Thus, the domain that dictates HD oligomerization resides solely in the OD, which produces a heptameric ring. Additionally, another significant difference between Rep and other SF3 helicases is the extent of conformational change that is triggered upon nucleotide binding. In Rep68, the AAA^+^ domain undergoes a large rigid body rotation with respect to the apo structure of ~44° while in E1 and SV40-Tag, this rotation is of only 4.8° and 12.8° respectively (Supplementary figure S8). The magnitude of the conformational change in Rep proteins is in part due to the configuration of the region that links the OD to the AAA^+^ domain. In E1 and SV40-tag, this region consists of a long α-helix that folds into the AAA^+^ domain. In contrast, Rep proteins have a long linker with a small α-helix of only four residues that interact loosely with the AAA^+^ domain (Supplementary Figure S8). From a functional perspective, the accepted model of DNA translocation by SF3 helicases during DNA unwinding does not require a large conformational change of the entire AAA^+^ domain. Instead, alternating conformations of the PS1βH during ATP binding and hydrolysis are sufficient to drive the process. Therefore, the question remains as to the functional significance of the large conformational change in Rep proteins. A simple explanation is that both E1 and SV40 are mainly processive helicases that need to translocate on DNA efficiently while AAV LReps are used in multiple processes that involved DNA structural reconfiguration and melting. In this context, the large conformational change seen in Rep68 may induce DNA distortions leading eventually to its melting. The degree of conformational change is similar in both heptameric and hexameric complexes and also involves the generation of a new DNA contact point at the PS1βH region. As a result, Rep68 ring structures contact DNA inside the channel at two locations separated by ~ 4-5 bp, a property that again is not shared other SF3 helicases, which contact DNA using only residues in the PS1βH region (31). The large conformational change may also explain how a mostly monomeric Rep40 may be able to unwind DNA without being a processive ring helicase, as the conformational change may be sufficient to unwind DNA without the need of forming a ring. Additional research will be needed to explore this hypothesis.

While the majority of AAA^+^ proteins, including most SF3 helicases, form stable oligomers irrespective of nucleotide binding, a small fraction of their members display a dynamic oligomeric behavior in a concentration and nucleotide-dependent manner (33). Our results show that Rep68 HD is of the latter type as its oligomerization is highly dependent on concentration, presence of nucleotide and DNA substrate. Moreover, the Rep HD is unique among SF3 family members by forming heptameric and hexameric rings. However, this property is not exclusive to AAV LReps, as some SF4 helicases have been reported as having this capability. For instance, gene 4 from bacteriophage T7 and the mitochondrial replicative helicase Twinkle also form mixtures of hexamers/heptamers (34–37). Compared to other helicases, what advantage does this property bring to AAV Rep proteins? Mechanistically, this ability might be a reflection of the multifunctionality of Rep proteins that are required to participate in multiple DNA transactions during the AAV life cycle, including DNA replication, transcription regulation, and genome packaging. Nevertheless, a common step in all these processes remains the assembly of a Rep complex after recruitment to the RBS site. In this context, the transition from heptameric to hexameric rings may be significant during the loading of Rep proteins onto dsDNA. Generally, the loading of a replicative ring helicase onto DNA requires the help of a helicase loader to open up the ring, thus allowing access to DNA. However, it has been hypothesized that in some helicase systems, an already open heptameric ring can easily assemble to either single or double-stranded DNA without the need of an additional factor (38–42). Once loaded, it switches to a processive hexameric form through the loss of one of the subunits in a process that is NTP and/or DNA dependent (43). A similar process seems fitting for Rep mediated processes. We have previously shown that Rep68 forms heptameric rings when bound to an AAVS1 dsDNA site (17). In addition, during AAV DNA replication, a necessary step during the terminal resolution reaction is the recognition of an RBS site, the formation of a Rep-DNA complex, and the eventual melting of the dsDNA. Based on our studies, we propose a mechanism where, after recruitment and binding to the RBS site, the LReps assemble initially as a heptameric complex (Figure 9). As a heptamer, the diameter of the central channel is wide enough to accommodate a dsDNA molecule, while in the hexameric form, this is not possible due to compression of the diameter of the channel to ~ 13 Å (Figure 7,8). Could DNA melting occur during the transition to a hexameric ring? Our data show that DNA deformation occurs in the heptameric complex, which could induce further structural deformations leading to DNA melting. We hypothesize that the asymmetry of the heptameric complex is one of the drivers of DNA deformation as the unbalanced neutralization of charges of the phosphate backbone is a well-known cause of DNA bending (44). Furthermore, the presence of two contact points in the DNA and the large conformational change upon nucleotide binding can produce additional deformation and DNA melting. While future studies will be needed to verify this model, our data has revealed important insights into the mode of action of AAV Rep proteins.

**Figure 9.**
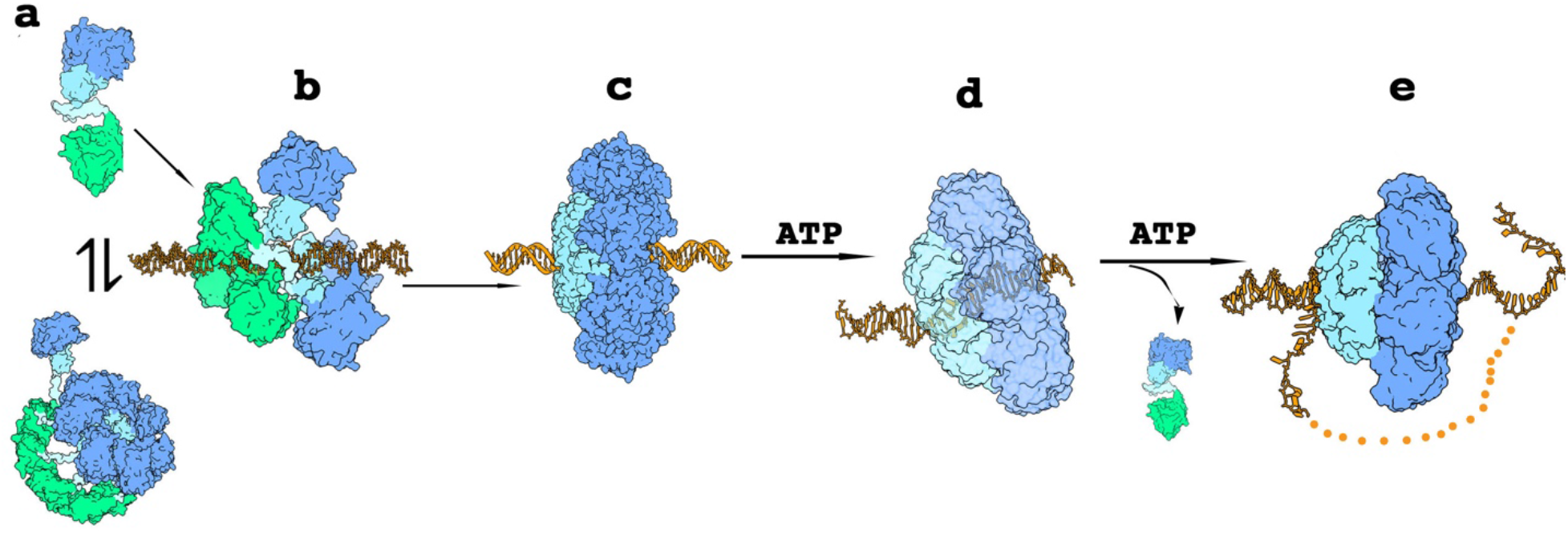
Model for assembly and DNA melting on dsDNA by AAV Re proteins. (a) Rep68 is in equilibrium between monomers and higher order oligomers including octamers. (b) Binding to RBS containing DNA sites is directed by OBD that interacts specifically with GCTC repeats and induces initial assembly. (c) A heptameric ring assembles around DNA and (d) upon binding ATP generates a deformation that produces strand separation. (e) Generation of ssDNA leads to a conversion from heptamer to hexamer with the expulsion of a Rep monomer. The OBD domains are not shown is steps (b) to (d) for simplification.

**Table 2.**
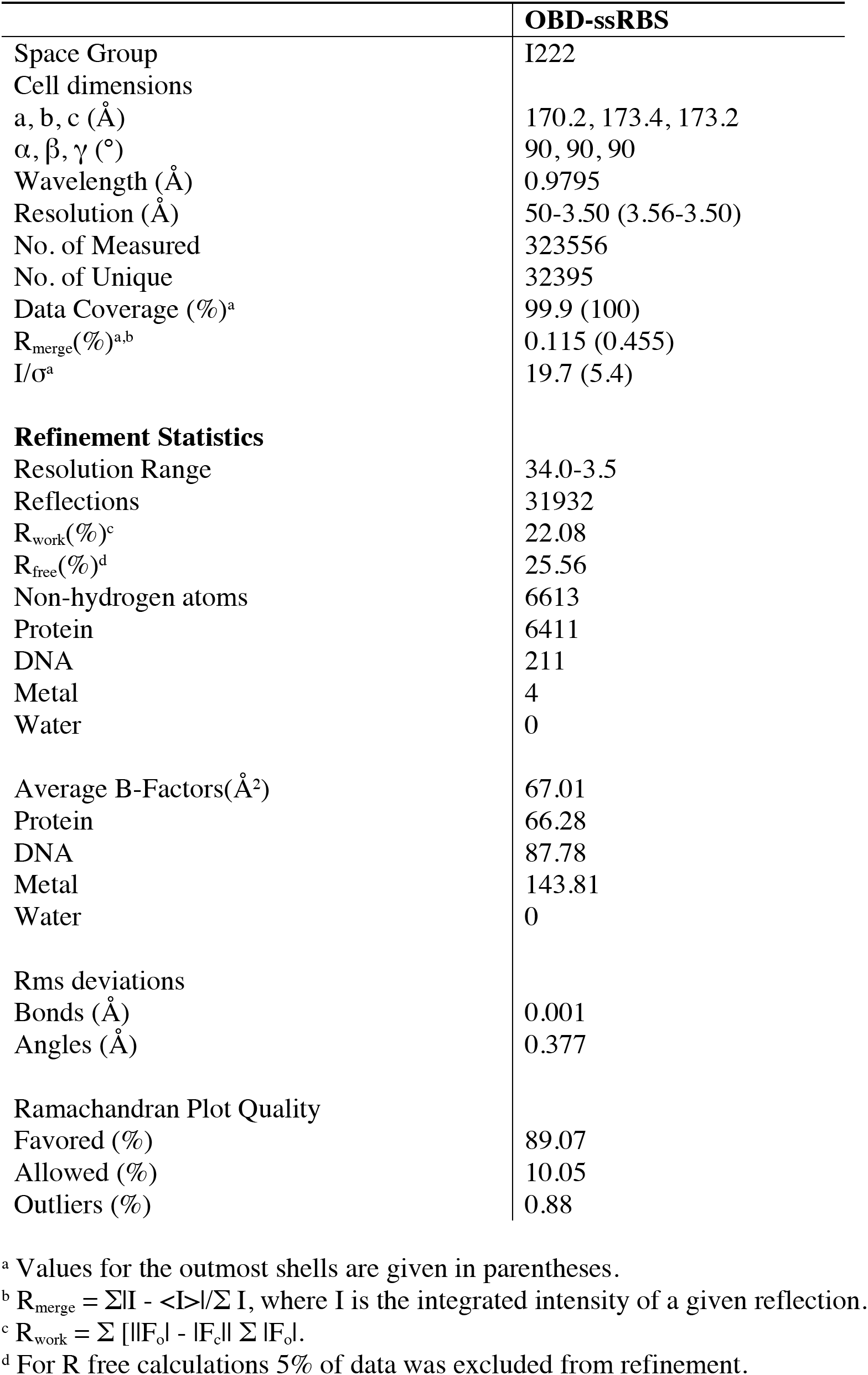
X-ray Data Collection and Refinement Statistics.

## Author Contributions

V.S., E.H., C.R.E., designed research; V.S., R.J., F.M., B.V., FZP, KS, C.D., M.E., K.D., and C.R.E. performed research; V.S., and C.R.E. wrote the paper with all authors providing comments or revisions.

## Acknowledgments

We would like to thank Ed Eng, Venkata Dandey, Elina Kopylov and outstanding staff from the National Center for Cryo-EM access and Training (NCCAT). We would like to acknowledge Lauren Hales Beck, Nancy Meyer from the Pacific Northwest Cryo-EM Center (PNCC) supported by NIH grant U24GM129547.

## Data availability

All cryo-EM densities and atomic coordinates have been deposited to EMDB. Densities for the DOC apo and ATPγS-bound structures are EMD-XXXX and EMD-XXXX respectively. Rep68-ssAAVS1-ATPγS density is EMD-XXX and PDB: XXXX. Rep68-ssdT_25_ hexamer density is EMD-XXX and PDB:XXXX. Xray structure coordinates for AAV2 OBD-ssRBS complex is PDB:XXXX.

## Funding

CRE is supported by National Institute of Health (NIH) R01 (GM124204).

EH is supported by a start-up grant from KU Leuven and research grant from the Flemish Research Council (FWO-G0C3220N).

A portion of this research was carried out at the NCCAT and the Simons Electron Microscopy Center located at the New York Structural Biology Center, supported by the NIH Commond Fund Transformative High Resolution Cryo-Electron Microscopy program (U24 GM1 129539) and by grants from the Simons Foundation (SF349247) and NY State. A portion of this research was supported by NIH grant U24GM129547 and performed at the PNCC at OHSU and accessed through EMSL (grid.436923.9), a DOE Office of Science User Facility sponsored by the Office of Biological and Environmental Research.

### Materials and Methods

#### Cloning and Mutagenesis of Rep Expression Constructs

All mutant proteins were generated using the pHisRep68/15b plasmid which contains the AAV2 Rep68 ORF (1-536) or the truncated form (1-490) were subcloned in vector PET-15b (Novagen). Site-directed mutagenesis for all mutants was generated using the QuickChange® mutagenesis kit (Stratagene). The sequences of all constructs were confirmed by DNA sequencing (GeneWiz).

#### Protein expression and purification

All proteins were expressed using the pET-15b vector, expressed in E. coli BL21(DE3) cells (Novagen), and purified as described before (23). The final buffer contains (25 mM Tris-HCl [pH 8.0], 200 mM NaCl, and 2mM TCEP). His6-PreScission Protease (PP) was expressed in BL21(DE3)-pLysS at 37 °C for 3 h, in LB medium containing 1 mM IPTG. Cell pellets were lysed in Ni-Buffer A (20 mM Tris-HCl [pH 7.9 at 4 °C], 500 mM NaCl, 5 mM Imidazole, 10% glycerol, and 1 mM TCEP). After five 10-s cycles of sonication, the fusion protein was purified using a Ni-column – equilibrated in Ni-buffer A. Protein eluted was desalted using buffer A and a HiPrep™ 26/10 desalting column (GE Healthcare). His-PP tag was removed by PreScission protease treatment using 150 μg PP /mg His-PP-Rep68. After overnight incubation at 4 °C, buffer was exchanged using the same desalting column and Ni-Buffer A. Subsequent Ni-column chromatography using the buffer B (same as buffer A but with 1 M imidazole), was performed to remove the uncleaved fusion protein, and untagged Rep68 was eluted with 30 mM imidazole. Rep68 was finally purified by gel filtration chromatography using a HiLoad Superdex 200 16/600 PG column (GE Healthcare) and Size Exclusion buffer. N-terminus His6-tagged WT and mutant Rep68 proteins were concentrated to 2 mg/ml, flash-frozen in liquid N_2_, and kept at −80°C until use.

#### Gel-filtration experiments

Gel-filtration studies were performed with the Rep68* concentration held constant at 30 μM and the DNA held constant at 10 μM. The gel-filtration buffer used was: 10 mM Na(PO_4_)_2_ pH 7.0, 150 mM NaCl 1 mM TCEP and all results were plotted using Origin (Origin labs). The experiments shown in Supplementary Figure 2 were conducted on a Superose 6 10/300 GL column (GE Healthcare) while the experiment shown in Figure 2 was conducted on a Superose 6 Increase 10/300 gel filtration column (GE Healthcare).

#### AAV Infectious particles assay

Two parallel systems were used. In one, AAVpro® 293T (Takara) cells were transfected in duplicates using CalPhos method with an AAV2 ITR-containing plasmid including GFP gene (pTRUF11), a plasmid expressing AAV2 Rep WT or mutants (pRC2) and Cap, and third plasmid containing adenovirus helper protein (Takara). 48 hours post transfection, cells were harvested and lysed using AAV extraction solution A & B (Takara) as per manufacturer’s recommendation. HT1080 cells were infected with increasing amount of viral solution and percentage of GFP-positive cells was determined after three days of infection. In the second protocol, HEK293T cells were triple transfected in a 6-well format using polyethylenimine (PEI) Max with 2 μg of an AAV2 ITR-containing plasmid including GFP as transgene, 1.6 μg of a helper plasmid expressing AAV2 Rep (WT or mutant) and Cap, and 1.6 μg of a third construct containing the adenovirus helper functions (pXX6, University of North Carolina Vector Core Facility). 72h post-transfection, 1 mL of the supernatant was spun down to clear the cellular debris and increasing volumes of supernatant were used to transduce HeLa cells at 60-70 % confluency. Before transduction, the medium of the HeLa cells was replaced with DMEM + 5 % FCS. 48h after transduction, the percentage of GFP-positive HeLa cells was determined by flow cytometry (Accuri™ C6 Plus; BD Biosciences) (26).

#### Analytical ultracentrifugation

Sedimentation velocity experiments were carried out using a Beckman Optima XL-I analytical ultracentrifuge (Beckman Coulter Inc.) equipped with a four and eight-position AN-60Ti rotor. Rep protein samples were loaded in the cells, using in all cases buffer used in the final purification step. Samples in double sector cells were centrifuged at 25,000 rpm for Rep68 proteins. For OBD and linker constructs sedimentation was performed at 40,000 rpm. In all experiments, temperature was kept at 20 °C. Sedimentation profiles were recorded using UV absorption (280 nm) and interference scanning optics. For the analysis of the results the program Sedfit was used to calculate sedimentation coefficient distribution profiles using the Lamm (45).

#### Helicase Assay

The substrate used in this assay is a heteroduplex DNA consisting of an 18-bp duplex region with a 10-nucleotide 3’ tail at the bottom strand. The top strand (trap-DNA) is labelled at the 5’ end with fluorescein and is released upon unwinding. The sequences used are 5’-F-CATATGGAGCAGAACAGA-3’ for the trap DNA and 5’-AGACAAGACGAGGTATACAAAAAAAAAA-3’ for the complementary strand. All reactions were performed in a buffer containing 25mM HEPES, 50mM NaCl (pH 7.0) at a total volume of 50μl. 1mM of protein was mixed with 0.5mM of double stranded F-DNA (18ADT10A) and 2.5μM of single stranded DNA (18s), and then added to the mix of buffer described above containing 5mM of both ATP and MgCl2. Reaction was incubated at 25°C for 1 hr. EDTA was used to stop the reaction at a final concentration of 20μM. Aliquots of 10μl were loaded in a 12% bis-acrylamide gel (30%) (19:1) using 6X-loading dye (0.25 xylene cyanol FF, 30% glycerol). For the densitometry and analysis of the bands, a Gel Doc EZ Imager was used, together with the automatic lane and band detection tool. Lane background subtraction, white illumination and an activation time of 300 sec were used for the analysis.

#### EM sample preparation

The Rep68-dT_25_ complex was prepared as previously described (23). In short, Rep68* was mixed with a 25-mer poly-dT oligonucleotide (dT_25_), purified to homogeneity and concentrated in the presence of 0.05% n-octyl-β-D-glucopyranoside (OG). The Rep68-AAVS1-15 ssDNA complex was made by incubating 30μM protein with 15μM ss AAVS1-15 DNA in reaction buffer (10 mM Na(PO_4_)_2_ pH 7.0, 150 mM NaCl 1 mM TCEP) buffer for 15 minutes at room temperature. Complex was concentrated to 200μL using Amicon Ultra-4 centrifugal filter (Millipore) and loaded on Superose 6 Increase 10/300 gel filtration column (GE Healthcare) pre-equilibrated with reaction buffer. Eluted complex was concentrated back to 1/8^th^ and 1/16^th^ of original protein concentration. C-Flat carbon grids CF1.2/1.3-4C, 400 mesh Cu (Electron Microscopy Sciences CF413-50) were glow-discharged for 45 seconds with amylamine using a PELCO easiGlow™ glow-discharge system. Just before spotting the sample, 5mM MgCl_2_ and 5mM ATPγS (final concentration) was added to the complex and incubated at room temperature for 5 minutes. 3.5 μL sample was spotted, blotted and plunged froze into liquid ethane using Vitrobot unit and stored in liquid nitrogen.

#### Cryo-EM grid preparation and data collection

C-flat grids were glow-discharged for 40 seconds with amylamine using a PELCO easiGlow™ glow-discharge unit and spotted with 3.0 μL of a 0.5 mg/ml Rep68-dT25 sample. The sample was manually blotted for 1.5 seconds and plunged into liquid ethane. For storage, the samples were stored in liquid nitrogen. To obtain more side views, 0.05% octyl-beta-glucoside (OG) was added to the sample and frozen as previously described (23). Initial screening was done on a Tecnai F20 electron microscope equipped with a 4k × 4k ultrascan CCD camera and preliminary data collection on a Titan Krios at the University of Virginia Molecular Electron Microscopy core facility. Final data sets for the DOC were collected at the National Center for CryoEM Access and training (NCCAT) located at the New York Structural Biology Center on a Titan Krios in counting equipped with a Gatan K2 Summit direct detector. For the apo-data, 40 movie frames were recorded at 165,000 magnification and total dose of 92.02 electrons per Å^2^ at a pixel size of 0.85 Å. For the ATPγS complex, the samples were collected at 22,500x at a pixel size of 1.073 Å and total dose of 55.46 electrons per Å^2^. Data for the Rep68-ssAAVS1-ATPγS were collected at the Pacific Northwest Center for Cryo-EM (PNCC) on a Titan Krios equipped with a Falcon 3 direct detector. Samples were collected at 59,000x magnification at a pixel size of 1.415 Å and total dose of 50 electrons per Å^2^ (TABLE 1).

#### Cryo-EM image processing and model building

For all apo DOC complex, micrographs frames were aligned and superimposed using Motioncorr2 (46). Contrast transfer function (CTF) parameters were calculated with CTFfind4 and automated particle image selection were done in RELION (47, 48). Selected particles (118,531) were imported into cryoSPARC v.2.3 to generate reference-free 2D classification to remove false particles and junk classes. The selected particles (62,512) were extracted and used to generate initial models. C1 symmetry was applied during all refinement process. Two 3D models were further processed using heterologous refinement and a final model was processed using 46,031 particles using non-uniform refinement to an overall resolution of 4.6 Å using the 0.143 cutoff.

For the DOC-ATPγ data, all steps were performed using cryoSPARC starting with 4555 micrographs. Preprocessing was done using Motioncoor2 and CTFfind4. A set of 294,324 particles were visually inspected through a series of passes of 2D classification and 2D selection to remove junk and low-quality particles, resulting in 78,701 good particles. After generating an initial model, and 3D non-uniform refinement the final map had an overall resolution of 5.2 Å using the 0.143 cutoff.

For the Rep68-ssAAVS1 complex, all steps were performed in cryoSPARC starting with 2669 movies. The movie frames were aligned and summed using Motioncorr2 to obtain the final dose-weighted images. Estimation of the CTF was performed using CTFfind4. In total 907,616 particles were selected from 2584 images. 2D classification resulted in a mixture of hexameric and heptameric classes, that based on inspection of the top views correspond to about 52% and 48% respectively.

For the Hexameric Rep68-dT_25_ complex, grids were prepared with Rep68-dT_25_ complex as described previously. After DOC purification on a Superose 6 column, ATPγS was added to a final concentration of 5mM. 2D classes contain mixtures of DOC and hexameric particles with a total of 578,679 particles. 2D class selection of hexameric complex resulted in 61,195 particles that were used for 3D reconstruction.

All molecular models were built with Chimera, Coot and refined with Phenix (49–51) using the local refined maps. Electrostatic surface potential calculation was done using Pymol (The PyMOL Molecular Graphics System, Version 2.0 Schrödinger, LLC). Crystal structure of OBD (PDB:5BYG) was used to build the double-octameric model. Initially, 16 individual OBDs were manually docked into the cryo-EM map and oriented using Fit in Map in Chimera. Resulting model of the double-octameric OBD was further refined using rigid body refinement in PHENIX. A similar approach was used to build the apo heptameric HD using the crystal structure of AAV-2 Rep40 (PDB:1S9H). For the ATPγS complex, during initial refinement in Phenix, two rigid bodies were defined for each helicase subunit. The rigid body 1 contained the OD (residues 1-277) and rigid body 2 the AAA^+^ domain (residues 278-490). All superposition analysis of the models was done using the program LSQMAN (52).

#### Crystallization and X-ray structure determination

The oligonucleotides used for crystallization were purchased from Integrated DNA Technologies, Inc. Site A: 5’-ATATGCTCGCTCTT-3’. DNA was purified on a MonoQ-5/50GL column. The purified DNA was, desalted, lyophilized, and resuspended in TE buffer (10mM Tris-HCl, pH 8.0, 40mM NaCl, 1mM EDTA). The oligonucleotides were mixed in 1:1 molar ratio, heated to 90°C for 5 minutes and cooled slowly to room temperature. OBD and double stranded DNA were mixed in a 1:1.1 molar ratio. Complex was concentrated to a final concentration of protein of about 20 mg/ml. The buffer concentration was exchanged during the concentration process to 25mM Tris-HCl, pH 7.5,100mM NaCl and 1 mM TCEP. Crystallization experiment was carried out using both hanging- and sitting-drop methods with commercially available screening kits at 4°C. Crystals grew from 3μl hanging drop after 2-3 weeks. The best crystals obtained from reservoir solution containing 10mM Na cacodylate, pH 6.5, 30% PEG400 and 0.2M LiCl. The crystals belonged to the space-group I222 with unit cell dimensions a=170.1 Å, b=173.4 Å, c=173.1 Å. Diffraction data were collected at the National Synchrotron Light Source (NSLS) at the Brookhaven National Laboratories beamline X6a. The data was processed with the program HKL2000 (53) and the structure was solved by molecular replacement using the program PHENIX (51). We used the structure of the AAV2 OBD as a search model (pdbid 4ZQ9). Model building was carried out using PHENIX and manual building using the program COOT (50).

---

**Table S1.** Cryo-EM data collection, refinement and validation statistics.

|  | Apo-(DOC) |  | ATPγS-(DOC) | Heptamer-ssAAVS1-ATPγS | Hexamer-dT<sub>25</sub>- ATPγS |
| --- | --- | --- | --- | --- | --- |
| Detector | Gatan K2 Summit |  | Gatan K2 Summit | Falcon 3 | Gatan K2 Summit |
| Magnification | 165,000 x |  | 22,500 x | 59,000 x | 22,500 x |
| Voltage (kV) | 300 |  | 300 | 300 | 300 |
| Electron exposure (e<sup>-</sup>/Å<sup>2</sup>) | 92.03 |  | 55.46 | 50.0 | 50.0 |
| Defocus range (μm) | -1.0 to -2.2 |  | -1.5 to -3.0 | -2.4 to -1.1 | -1.25 to -2.75 |
| Pixel size (Å) | 0.825 |  | 1.073 | 1.415 | 1.073 |
| Symmetry imposed | C1 |  | C1 | C1 | C1 |
| # images | 1449 |  | 4555 | 2669 | 1722 |
| Initial particles | 118,531 |  | 294,324 | 433,139 | 578679 |
| Final particles | 46,031 |  | 78,701 | 193,869 | 61195 |
| Map resolution (Å) | 4.6 |  | 5.2 | 4.4 | 5.0 |
| FSC Threshold | 0.143 |  | 0.143 | 0.143 | 0.143 |
| Refinement |  |  |  |  |  |
| Cryo-EM Model | OBD-DOC | HD1 | HD1 | HD-Heptamer | HD-Hexamer |
| Initial models used (PDB code) | 5BYG | 1S9H | 1S9H | 1S9H | 1S9H |
| Model resolution (Å) | 4.4 | 5.9 | 7 | 5.5 | 5.4 |
| FSC threshold | 0.143 | 0.143 | 0.143 | 0.143 | 0.143 |
| Model composition |  |  |  |  |  |
| Non-hydrogen atoms | 26392 | 15162 | 15043 | 15403 | 13014 |
| Protein residues | 3168 | 1904 | 1904 | 1904 | 1656 |
| DNA | 32 | - | - | - | - |
| B factors (Å<sup>2</sup>) |  |  |  |  | 134 |
| Protein | 156.5 | 69 | 43 | 43 | 134 |
| DNA | 22.6 | - | - | - | - |
| r.m.s.d |  |  |  |  |  |
| Bond lengths (Å) | 0.003 | 0.006 | 0.003 | 0.004 | 0.003 |
| Bond angles (°) | 0.893 | 1.5 | 0.665 | 0.661 | 0.920 |
| Validation |  |  |  |  |  |
| Molprobity score | 1.85 | 1.85 | 3.09 | 3.25 | 2.01 |
| Clash Score | 4.88 | 2.59 | 27.19 | 32.66 | 6.72 |
| Ramachandran plot (%) |  |  |  |  |  |
| Favored | 88.01 | 86.72 | 91.4 | 94.4 | 86.72 |
| Allowed | 10.97 | 12.38 | 8.6 | 5.22 | 12.38 |
| Outliers | 1.02 | 0.9 | 0 | 0.37 | 1.71 |

**Figure S1.**
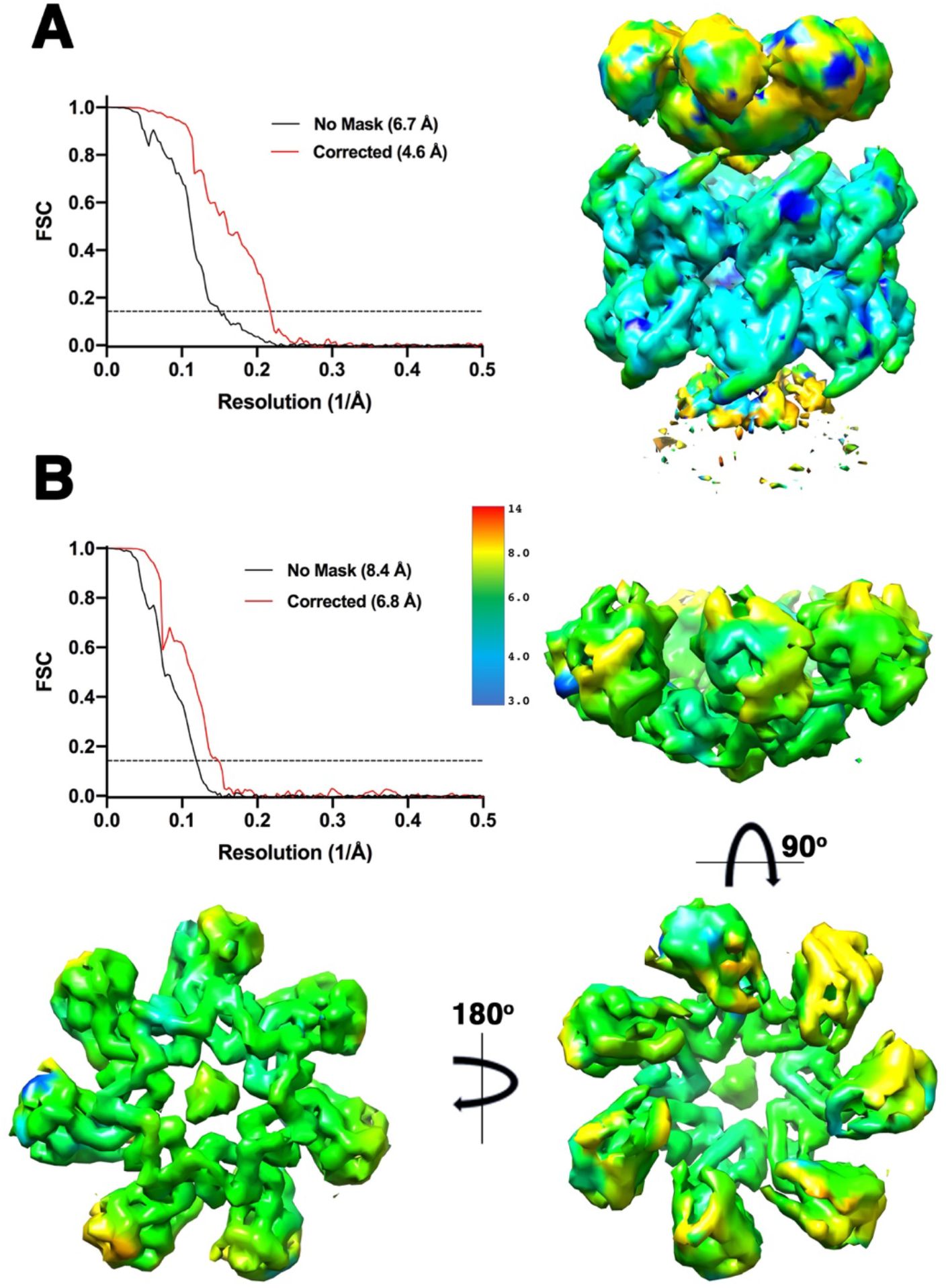
Resolution of Rep68-ssDNA DOC. (A) Fourier Shell Correlation (FSC) plot generated by cryosparc (left) and Cryo-EM map colored by resolution (right). (B) FSC plot of Helicase domain focused refinement (left) and three views of the HD1 cryo-EM map colored by resolution (right and bottom panels).

**Figure S2.**
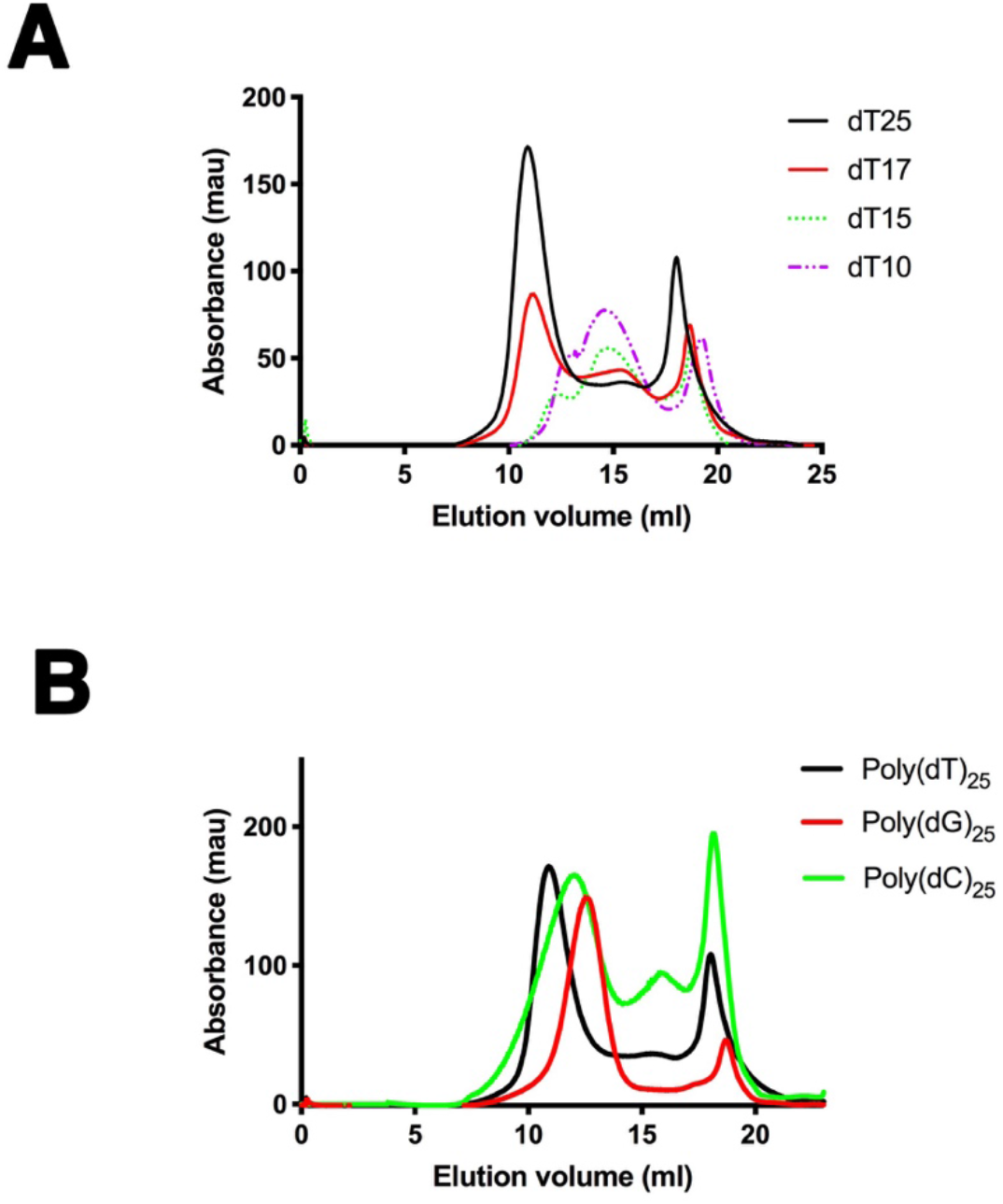
ssDNA structural determinants of DOC formation. (A) Rep68 (17 μM) was incubated with poly-dT ssDNA (2.8 μM) of different lengths and loaded on a Superose 6 10/300 GL column with flow rate of 0.5 ml/min. Protein elution was followed by UV detection at 280 nM. Elution profiles correspond to poly-dT 25-mer (black), 17-mer (red), 15-mer (green dotted) and 10-mer (purple dotted). (B) Elution profiles of Rep68 incubated with different ssDNA using conditions described in (A). Shown are the profiles of the incubation with poly-dT_25_ (black), poly-dG_25_ (red) and poly-dC_25_ (green).

**Figure S3.**
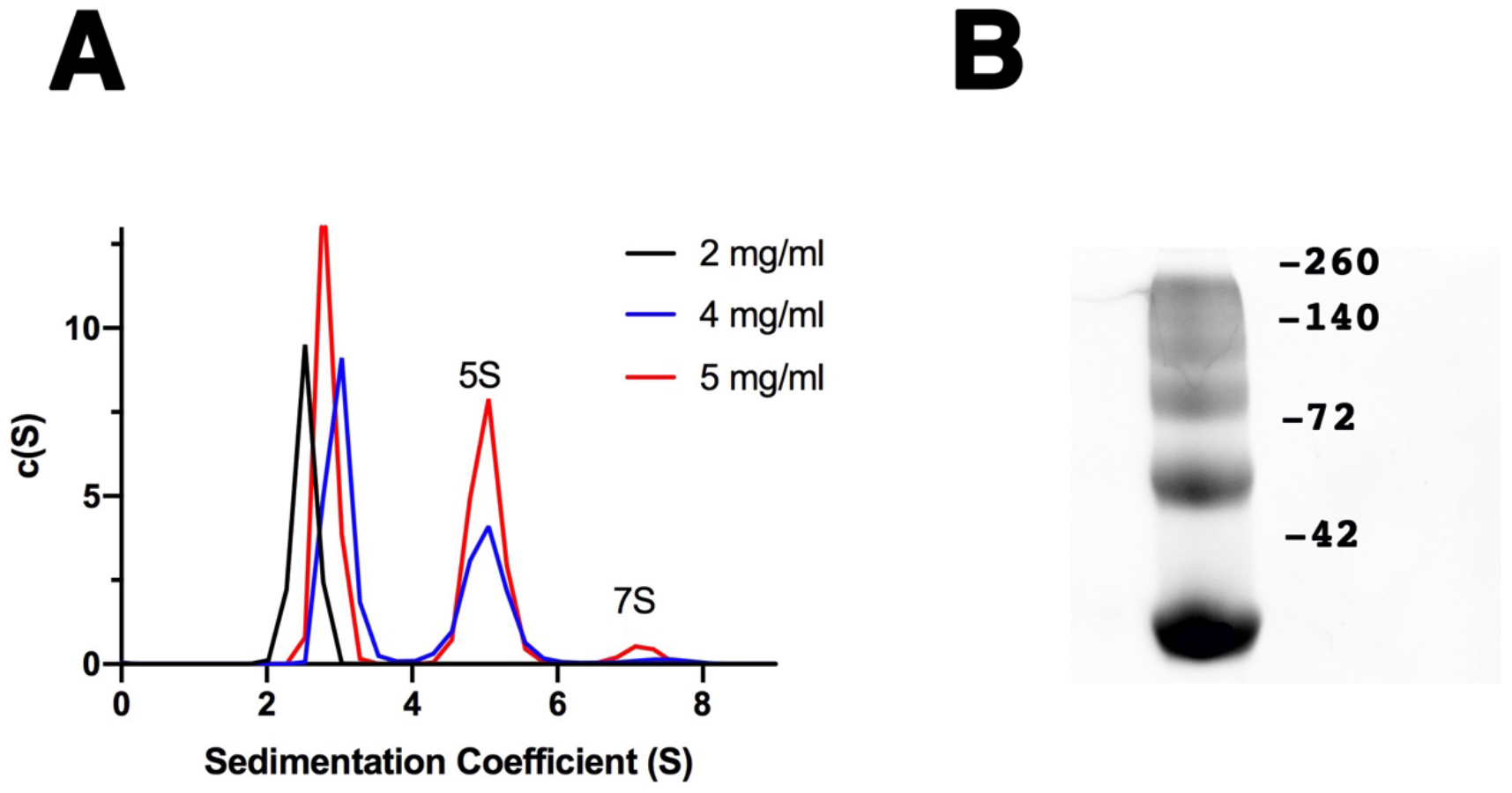
OBD forms octamers. (A) Sedimentation velocity data of OBD at 2mg/ml (black), 4 mg/ml (blue) and 5 mg/ml (red). Experiments were performed in Buffer A at 20°C on a Beckman XL-I analytical ultracentrifuge at 24,000 rpm. (B) Glutaraldehyde crosslinking of OBD at 5 mg/ml. Reactions were analyzed on a gradient SDS-PAGE (8-20%) stained-free gel and analyzed on a BioRAD GelDoc. Molecular weight of OBD (1-208) is 24.35 kDa. m,d,t,tt,o correspond to monomer, dimer, trimer, tetramer and octamer respectively.

**Figure S4.**
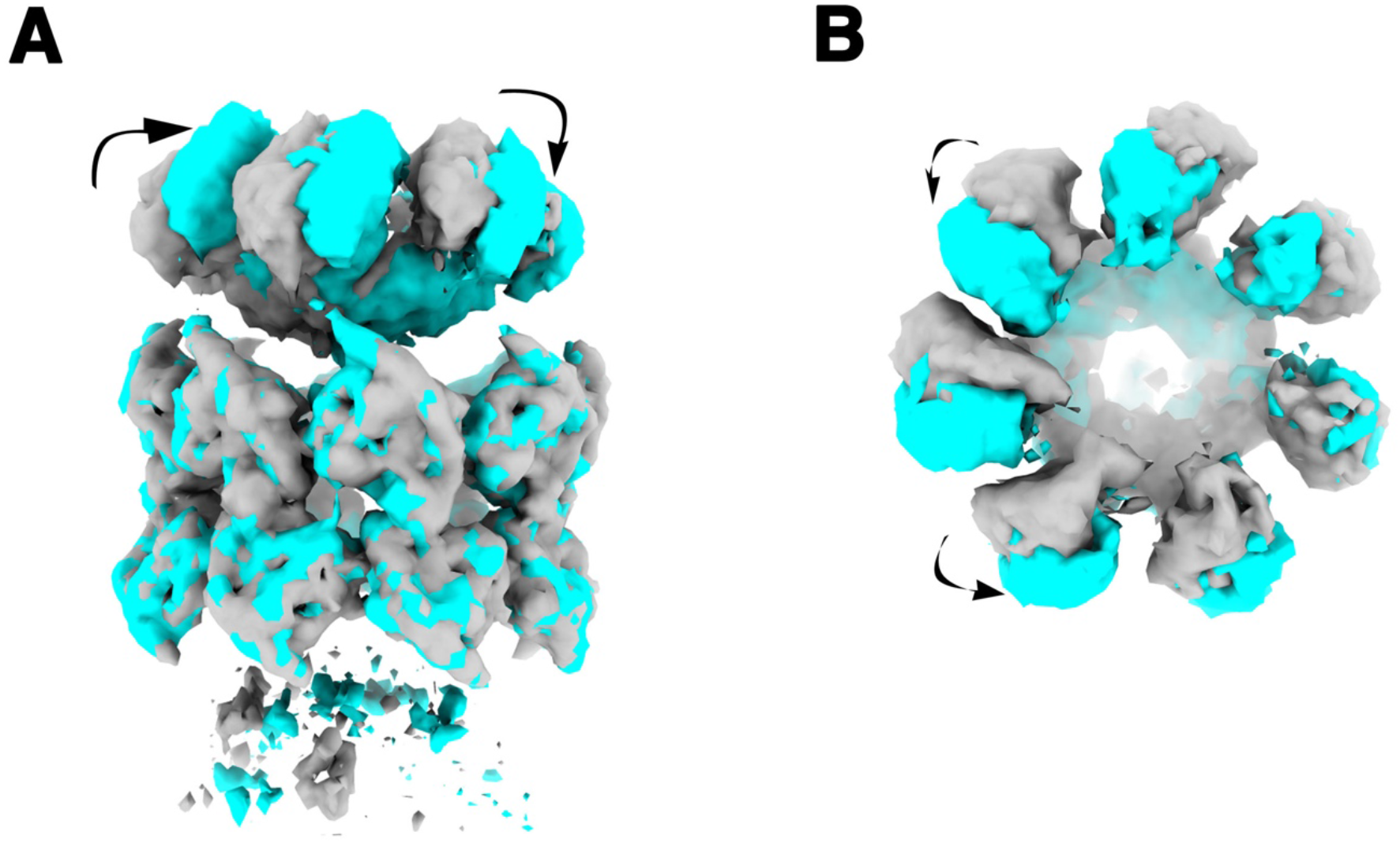
3D variability analysis of DOC. (A) and (B) Cryo-EM density showing two of the normal mode conformations in grey and blue. The arrows show the movement of the HD domains from one conformation to the other.

**Figure S5.**
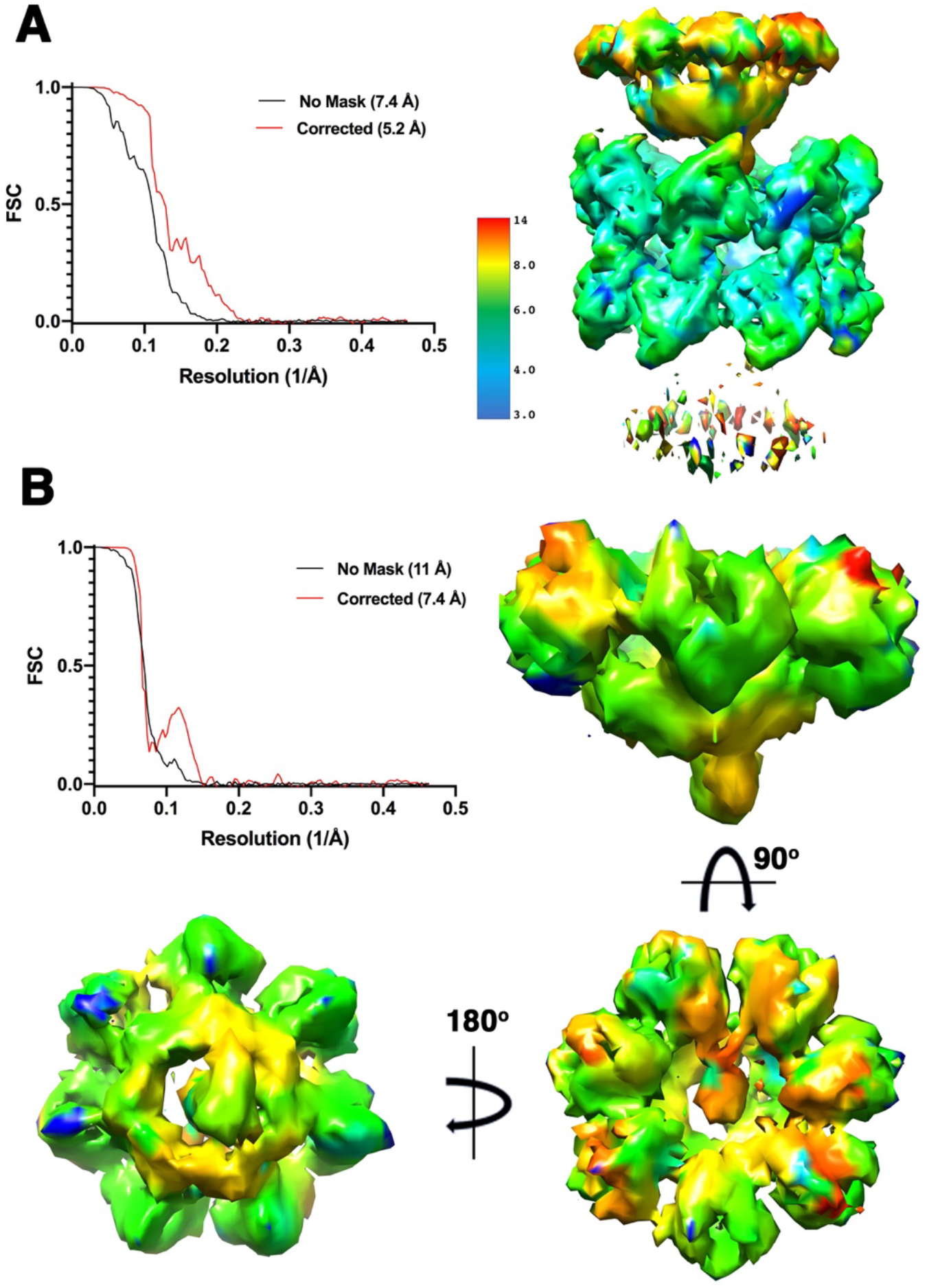
Resolution of Rep68-ssDNA-ATPγS DOC. (A) Fourier Shell Correlation (FSC) plot generated by cryosparc (left) and Cryo-EM map colored by resolution (right). (B) FSC plot of Helicase domain focused refinement (top left) and three views of the HD1 cryo-EM map colored by resolution.

**Figure S6.**
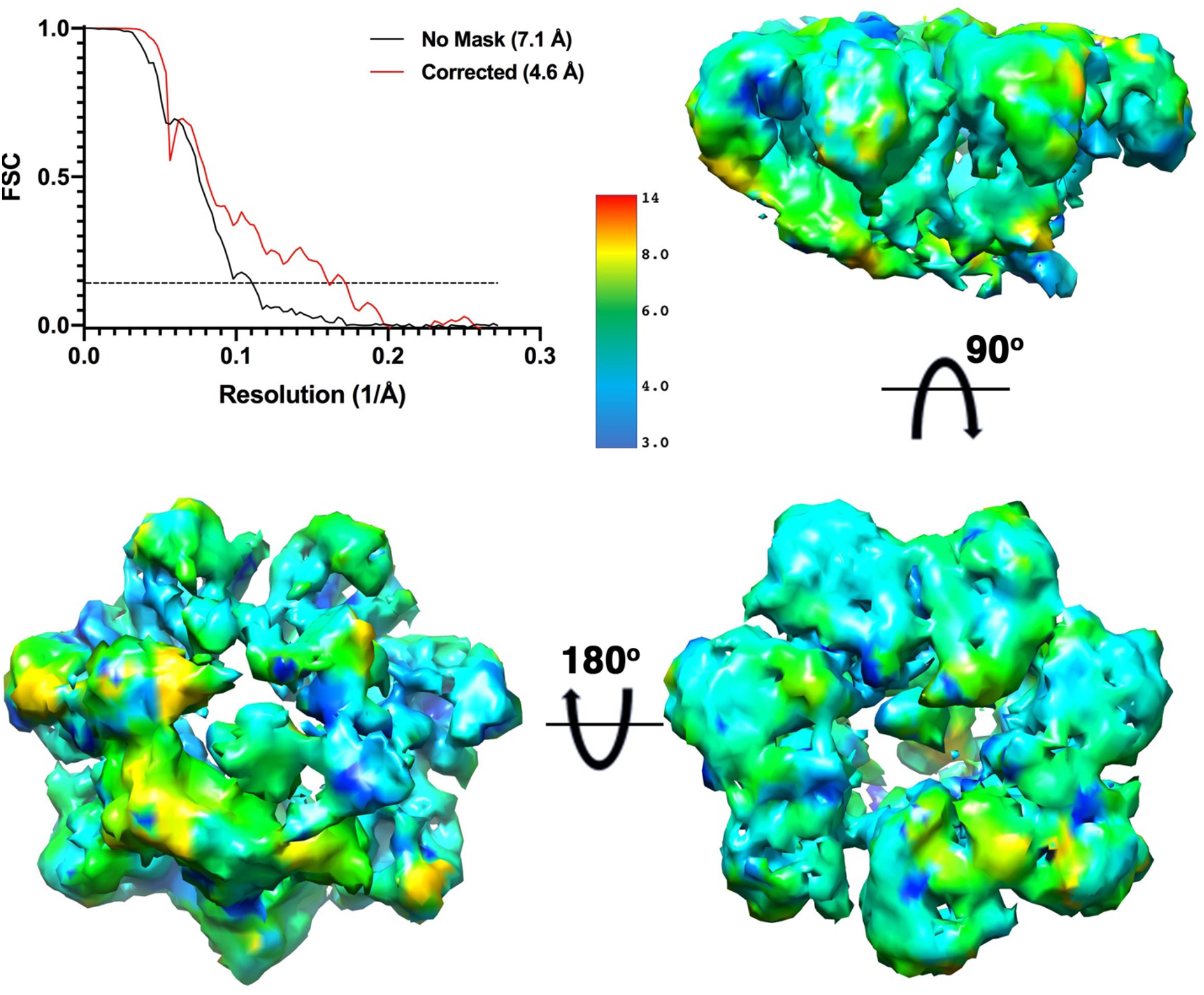
Resolution of Rep68-ssAAVS1-ATPγS heptameric complex. Fourier Shell Correlation (FSC) plot generated by cryosparc (left) and three views of the Cryo-EM focused refinement map colored by resolution (right and bottom).

**Figure S7.**
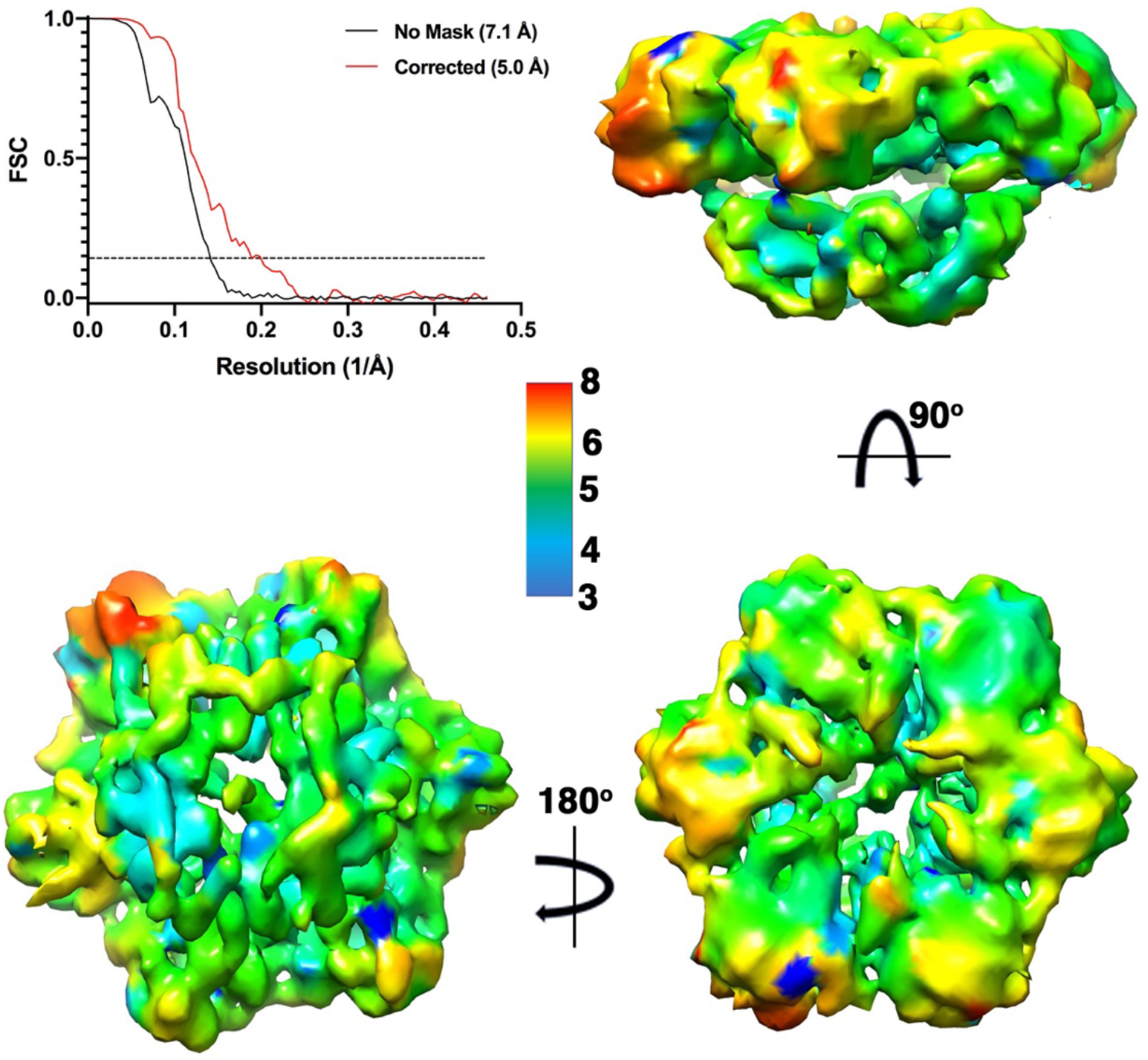
Resolution of Rep68-ssAAVS1-ATPγS hexameric complex. Fourier Shell Correlation (FSC) plot generated by cryosparc (left) and three views of the Cryo-EM focused refinement map colored by resolution (right and bottom).

**Figure S8.**
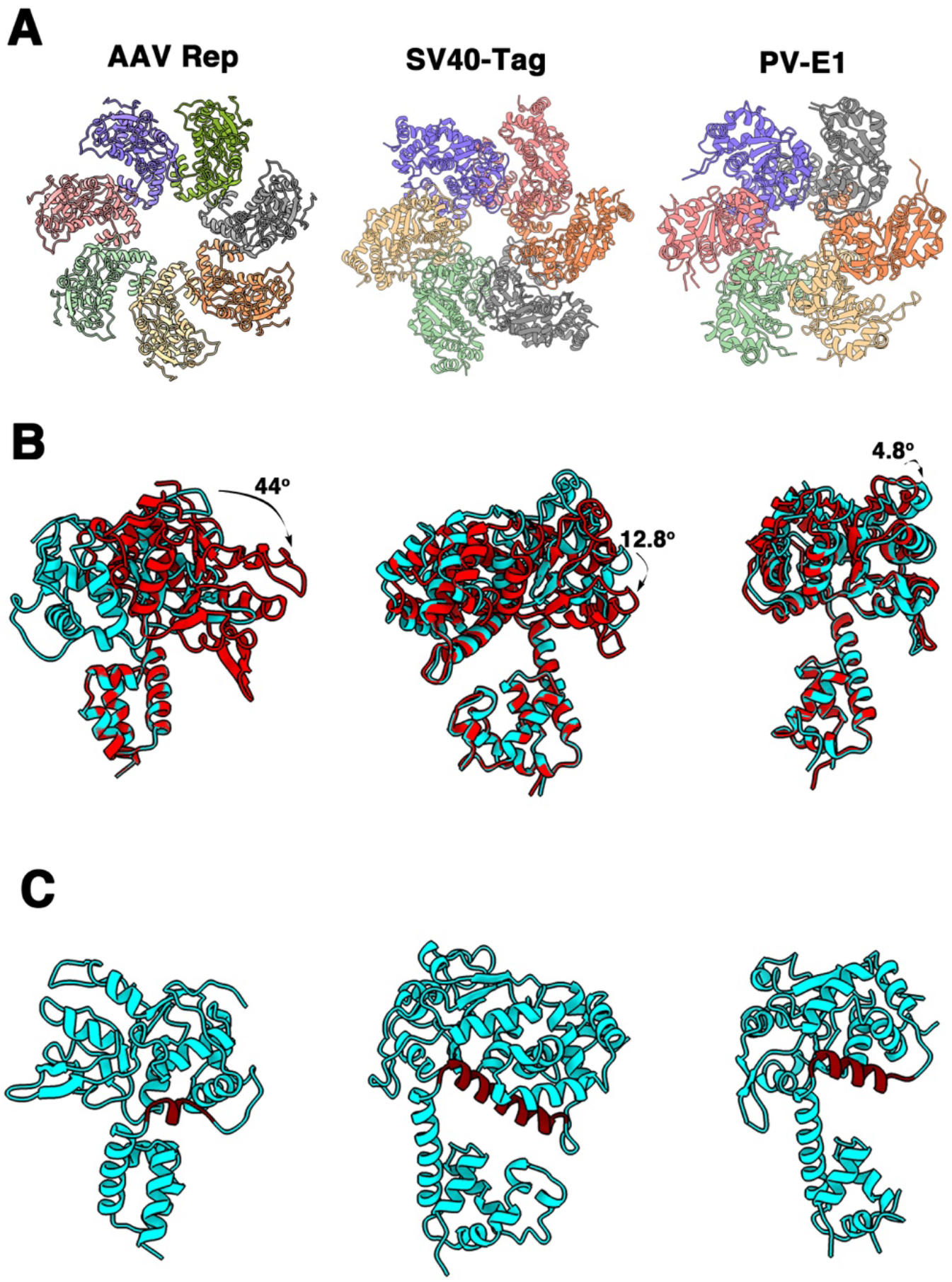
Comparison SF3 Helicases. (A) Ribbon representation of helicase domains of Apo structures of AAV Rep (left), SV40-Tag (center) and PV-E1 (right). (B) Comparison of rigid body movement of AAA^+^ domains of SF3 helicases upon nucleotide binding. (C) Comparison of region that connects OD to AAA^+^ domain in SF3 helicases. The regions are colored in maroon.

